# Genomic expression responses in *sensu stricto Saccharomyces* yeast to DNA damage induced by methyl methanesulfonate

**DOI:** 10.1101/2022.06.07.495055

**Authors:** Vignesh Ramachandran, Gregory Hatlestad, Travis White

## Abstract

**Background:** One way single-celled eukaryotes respond to DNA damage stress is by modifying their gene expression, facilitating genomic repair. Gene expression responses to DNA damage induced by methyl methanesulfonate (MMS) have been studied in the model organism *Saccharomyces cerevisiae*. However, lacking are investigations of the MMS stress responses in evolutionarily-related *sensu stricto Saccharomyces* species, including *Saccharomyces cerevisiae*.

**Methods:** Next-generation Illumina RNA-sequencing was to characterize the entire transcriptomes of four evolutionarily-related species of yeast, *S. cerevisiae*, *S. paradoxus*, *S. mikatae*, and *S. bayanus*, under control and experimental (MMS) conditions. Subsequent genomic studies included gene set enrichment analysis, promoter analysis, and concentration gradient studies.

**Results:** *S. mikitae and S. paradoxus* grew well in light of MMS while *S. bayanus* showed no growth. While there was fair overlap in induced and repressed genes, overall each species had unique expression responses. *S. paradoxus* and *S. bayanus* showed the most distinct changes with the former greatly inhibiting a large segment of its genome while the latter induced such segments. Gene set enrichment analysis revealed significantly modulated biologic, cellular, and molecular processes in each species. Promoter analysis revealed sets of induced/repressed transcription factors for genes highly modulated in the stress response. Concentration gradient studies of *S. cerevisiae* showed linear increase in gene expression of RAD54, DIN7, and IRC19 in response to increasing concentrations of MMS.

**Conclusion:** Overall, we depict the transcriptome changes of four evolutionarily-related *sensu stricto* yeast species and several functional genomic analyses to provide a novel understanding of their responses to MMS.

## Introduction

It is essential for organisms to maintain the integrity of their genomic material in order to survive and propagate. DNA damage may compromise the genetic material and cause genomic instability (Banin, Moyal et al., 1998). This may occur as a result of environmental stresses experienced during normal cell growth (Mager and De Kruijff, 1995; Ruis and Schuller, 1995; Gasch et al., 2000). In response to such environmental challenges, cells have developed intricate mechanisms of surveying for DNA damage and genomic stability during cell-cycle progression. This allows for complex responses to be mounted to ensure accurate propagation of genetic material (Hartwell and Kastan, 1994; Elledge, 1996).

Several cellular mechanisms to ensure genomic integrity during the cell cycle have been identified. Specifically, a response to DNA damage may be facilitated by cell-cycle arrest, alterations in gene expression, DNA damage repair, and cell death (Gasch et al., 2001; Branzei and Foiani, 2008). Maintenance of a stable genome is of particular importance to single-celled eukaryotic organisms as their sole cells are the basis of their existence. In yeast, altered transcription levels of genes in response to a stress condition have been previously implicated (Mager and De Kruijff, 1995; Ruis and Schuller, 1995; Morgan and Banks, 1997; Morano et al., 1998; McDaniel et al., 2017).

One method of studying gene expression changes in response to stresses in eukaryotes is by utilizing model organisms, such as *Saccharomyces cerevisiae.* Gene expression changes in response to environmental stresses in *S. cerevisiae* show significant and unique alterations for different stress conditions (Gasch et al., 2000). This is theorized to be due to the notion that the optimal response to a particular environmental stress is unique and not adequate for other stresses (Gasch et al., 2000; Rokhlenko et al., 2007). Specifically, the expression response of *S. cerevisiae* and the role of its yeast ATR homolog Mec1p in response to the DNA-damaging agents MMS and ionizing radiation have been studied in wild type and mutant cells. The results indicate that a broad and significant change in genomic expression is required by *S. cerevisiae* to cope with these stresses (Gasch et al., 2000).

*S. cerevisiae* is a part of the *sensu stricto* group, which includes other yeast species such as *S. paradoxus*, *S. mikatae*, and *S.bayanus* (Liti et al., 2013). Other studies report that as the evolutionary divergence between species of yeast increases, so too does their gene expression (Scannell et al., 2011). However, how the expression patterns of these evolutionarily-related species of yeast vary in response to stresses are not clearly elucidated in the literature. In this study, we depict the transcriptome changes of the four evolutionarily-related yeast *sensu stricto* yeast species and analyze the results through several functional genomic methods to provide a novel understanding of the unique yet also similar genome-wide expression patterns in response to MMS stress.

## Material and Methods

### Strains and Growth Conditions

**Table 1.** Yeast strains used in this study.

| Name | Genotype |
| --- | --- |
| *Saccharomyces cerevisiae* | wild type |
| *Saccharomyces paradoxus* | wild type |
| *Saccharomyces mikatae* | wild type |
| *Saccharomyces bayanus* | wild type |

### Differential Growth Characterizations

YPD plates were used for the control growth condition (Sherman, 2002) and YPD plates with 0.01% MMS by volume were used for the MMS experimental condition. Single colonies of each species of approximately similar size were used to inoculate four test tubes. The four liquid cultures were grown overnight in their individual test tubes with YPD media for approximately 16 hours. The cultures were then diluted to OD_600_ 0.8. Samples from the tubes were spotted via pipettes onto the YPD and MMS infused YPD plates in serial dilutions of 1/1, 1/10, 1/100, and 1/1000. Nanopure water was used as the diluting agent. Growth characterization was carried out for 72 hours.

### Overnight Sample Growth, Cell Lysis, and Total RNA Isolation

Single colonies of each of the four cultured yeast species from YPD plates were introduced into four separate test tubes containing liquid YPD media. The cultures were grown overnight for 16 hours and subsequently diluted to OD_600_ 0.2. The cultures were allowed to grow for one doubling period to OD_600_ 0.4 when each of the four cultures was split into two separate cultures of equal volume. One of the two cultures for each species had 0.03% MMS by volume introduced. Since the effects of MMS primarily manifest themselves during DNA replication, a doubling period was allowed to occur until OD_600_ 0.8. The OD_600_ values were confirmed by spectrophotometer. Cells were then collected by centrifugation at 4000 rpm for 5 minutes. The YPD above the cell pellet was decanted and the remaining cell pellets were stored at -20^०^C until RNA preparation. Total RNA extraction was subsequently performed using a modified procedure of (Schmitt et al., 1990). After completing the procedure, the RNA pellet was resuspended in 50 µl of nanopure water and stored at -20^०^C.

### Nanodrop, Gel Electrophoresis, and Quantitative PCR (qPCR)

The RNA samples extracted were tested for adequate concentration, integrity, and lack of ethanol contamination via Nanodrop spectrophotometric analysis. Gel electrophoresis was then used to visualize the samples for assurance that the samples were suitable for quantitative RT-PCR (qPCR) (Fleige and Pfaffl, 2006). Then, cDNA was created from a portion of the *S. cerevisiae* RNA sample. Verification that the stress response was induced was received by analyzing two genes known to be induced under MMS stress conditions in *S. cerevisiae*, RAD54 and DIN7 (Gasch et al., 2001). The normalizing gene ALG9 was used (Teste et al., 2009). The other three species did not have a study with which we could compare fold change values of RAD54 and DIN7 in response to 0.03% MMS stress conditions.

### Clustering and Java Treeview

Cluster analysis was performed using the program Cluster 3.0 (de Hoon et al., 2004), which utilizes the most commonly used clustering techniques to group genes according to similarities in gene expression. The filter of “% Present ≥ X” was applied to filter for genes that had data for at least 3 of the 4 species (75%). Hierarchical clustering settings included clustering of “Genes and Arrays”, “calculated weights”, and organization according to a similarity metric of “City-block distance”. The Clustering method utilized was “Average linkage”. The results were visualized using Java Treeview.

### Induced and Repressed Gene Overlap

In order to further examine the differences in expression patterns between the species at the gene level, Venn diagrams were created from separate lists of significantly induced and repressed genes (defined as fold changes ≥ 1 or ≤ -1).

### Gene Set Enrichment Analysis

Gene Set Enrichment Analysis (GSEA) is a computational technique used to determine whether previously defined sets of genes show statistically significant, concordant differences in two distinct biological states (i.e. control vs stress conditions). In our analysis, significantly regulated genes, defined for this analysis as fold changes that were ≥ +1 or ≤ -1, were inputted into WebGestalt (Wang, 2013) for each species of yeast. The settings used for the GSEA were as follows: “Enrichment Analysis”, “GO Slim Classification”, “scerevisiae_genome” (reference set for enrichment analysis), “hypergeometric” (statistical method), “BH” (multiple test adjustment), “0.05” (significance level), “2” (minimum number of genes in a category).

### Promoter Analysis

Yeastract Discoverer tools were used for motif extraction (Montiero et al., 2008). The Discoverer program utilizes structured motif discovery algorithms: MUSA (Motif finding using an UnSupervised Approach). We combined the lists of genes that were highly induced (≥ 3 folds) in the four yeast species. We repeated the same procedure for genes highly repressed (≤ -3 folds). The Discoverer program also identifies transcription factors associated with the motifs identified.

### Concentration Gradient Study

Yeast growth and RNA extraction were performed as previously described. The concentrations of MMS used for the stress conditions were: 0.06%, 0.05%, 0.04%, 0.03%, 0.02%, 0.01%, 0.001%, and 0% (control). The RNA was used to create cDNA to be used in qPCR as per the previously described procedure. The primer sets utilized were for the genes RAD54, DIN7, and IRC19. A primer set for the gene ALG9 was also used. ALG9 was used as the normalizing gene in this study (Teste et al., 2009).

## Results

### Differential Growth Characterizations

*S. paradoxus* and *S. mikitae* showed the greatest ability to grow in the MMS stress condition (Fig. 1). while *S. bayanus* demonstrated the least ability. In fact, *S. bayanus* had almost entirely no growth. The differing growth capabilities of these four species prompted investigation of their genome-wide expression responses to the MMS stress. This is grounded in the principle that the ability of an organism to adapt to stresses is due to changes in its expression patterns to maintain an internal environment conducive to proper cell function (Gasch et al., 2010).

**Figure 1.**
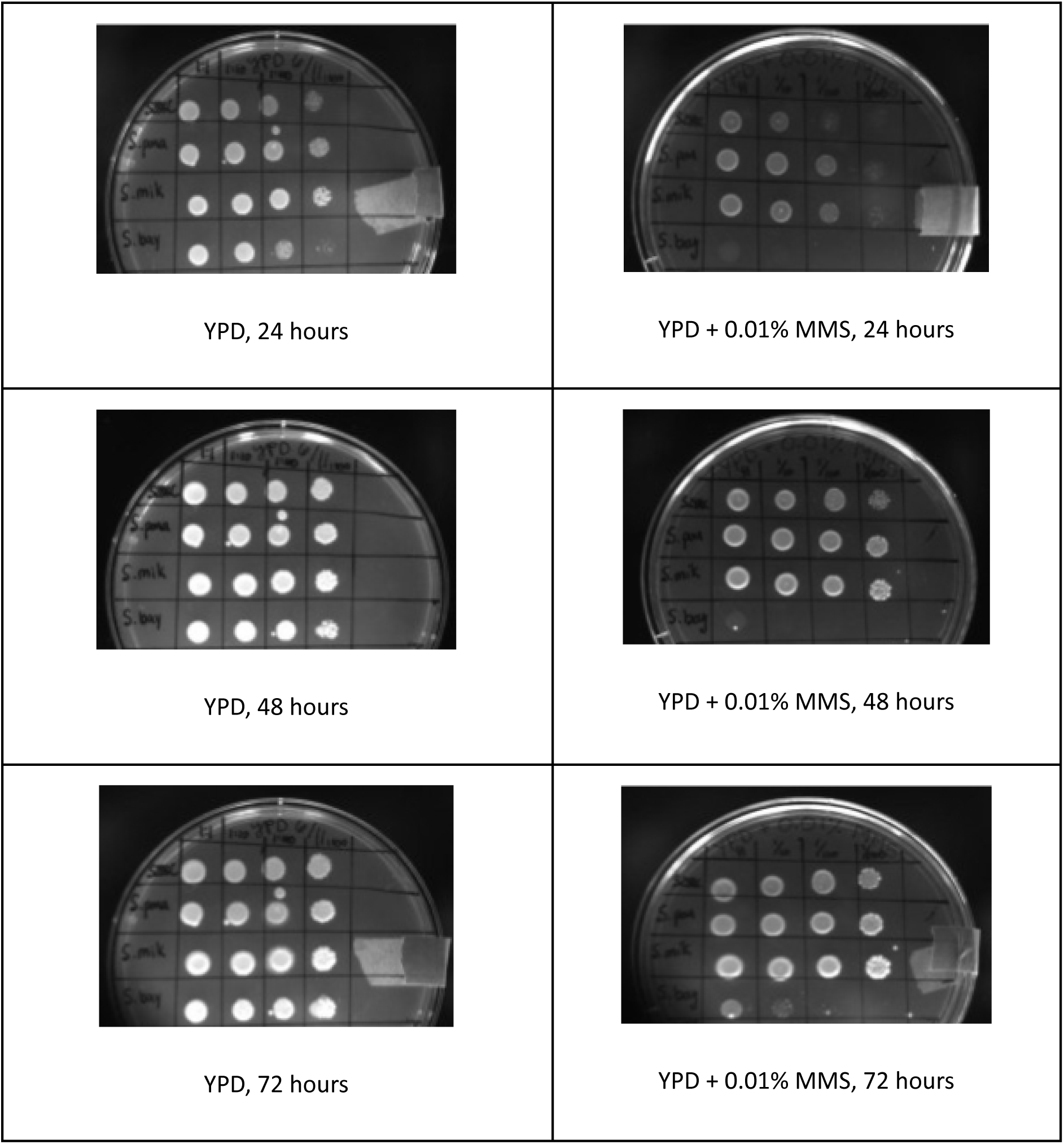
Differential growth characterizations of *S. cerevisiae*, *S. paradoxus*, *S. mikatae*, and *S. bayanus*, (top to bottom) with serial dilutions of 1/1, 1/10, 1/100, and 1/1000 (left to right).

### Next-Generation Illumina RNA Sequencing

Prior to conducting sequencing, quantitative polymerase chain reaction (qPCR) was utilized to examine if the stress was successfully introduced. Fold changes in the genes RAD54 (2.54) were compared to those in the literature (0.53 using 0.02% MMS) (Gasch et al., 2001). The results of the entire transcriptome changes in each species in response to MMS stress are depicted via heat maps generated in Java Treeview (Fig. 2). Red bands indicate induced genes and green bands indicate repressed genes. The level of induction or repression is indicated by the relative brightness of the bands. Brighter bands indicate more significant induction or repressed and vice versa. Grey regions indicate genes to which alignments were not made.

**Figure 2.**
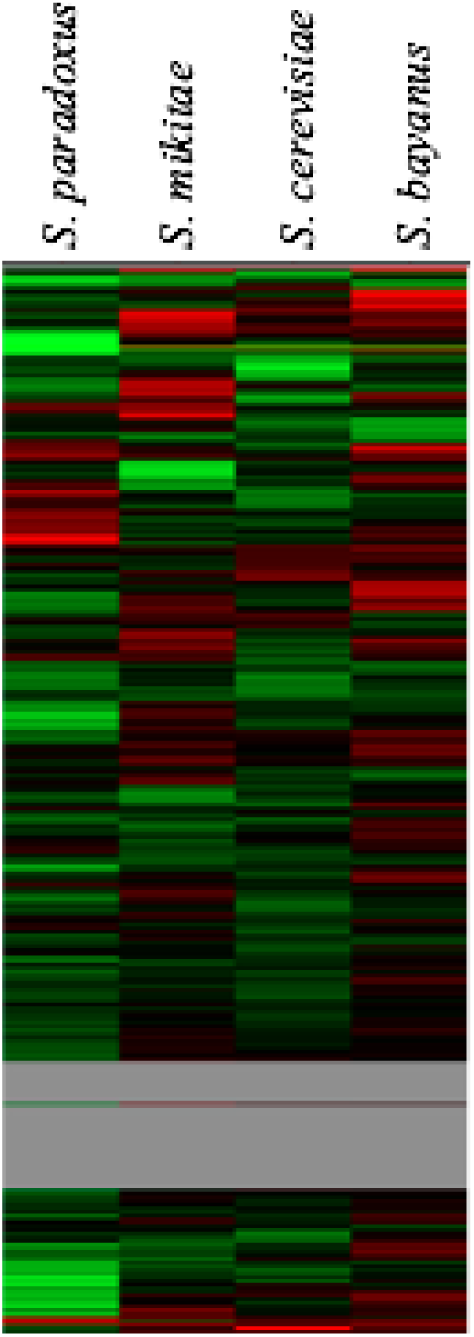
Visual representation of the transcriptome changes between control and experimental conditions for each yeast species. From left to right: *S. paradoxus*, *S. mikatae*, *S. cerevisiae*, *S. bayanus*.

Qualitative assessment of the overall expression patterns shows general similarity between *S. paradoxus*, *S. mikatae*, and *S. cerevisiae*. However, *S. bayanus* shows a different pattern of expression than those of the other three species in which a large section of the transcriptomes shows high levels of gene induction (Fig. 3). While *S. mikatae* also induces a subset of genes in this section, it is not as large or as significantly induced of a subset as that of *S. bayanus* displays. Additionally, *S. paradoxus* shows more overall gene repression and of a greater level than the other species (Fig. 4).

**Figure 3.**
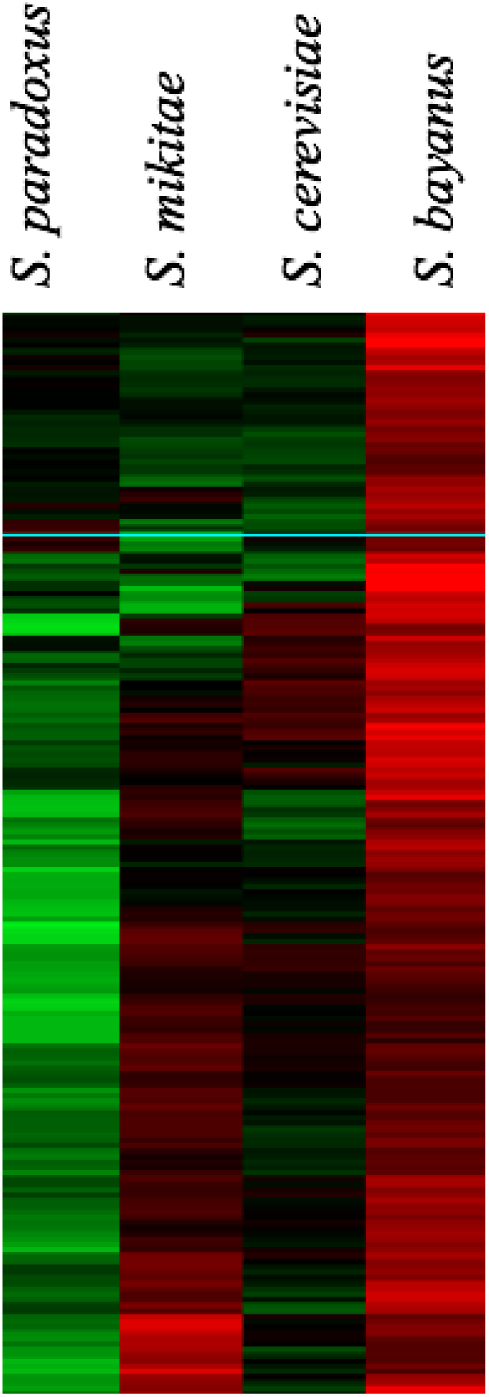
The primary region of induced genes in *S. bayanus*.

**Figure 4.**
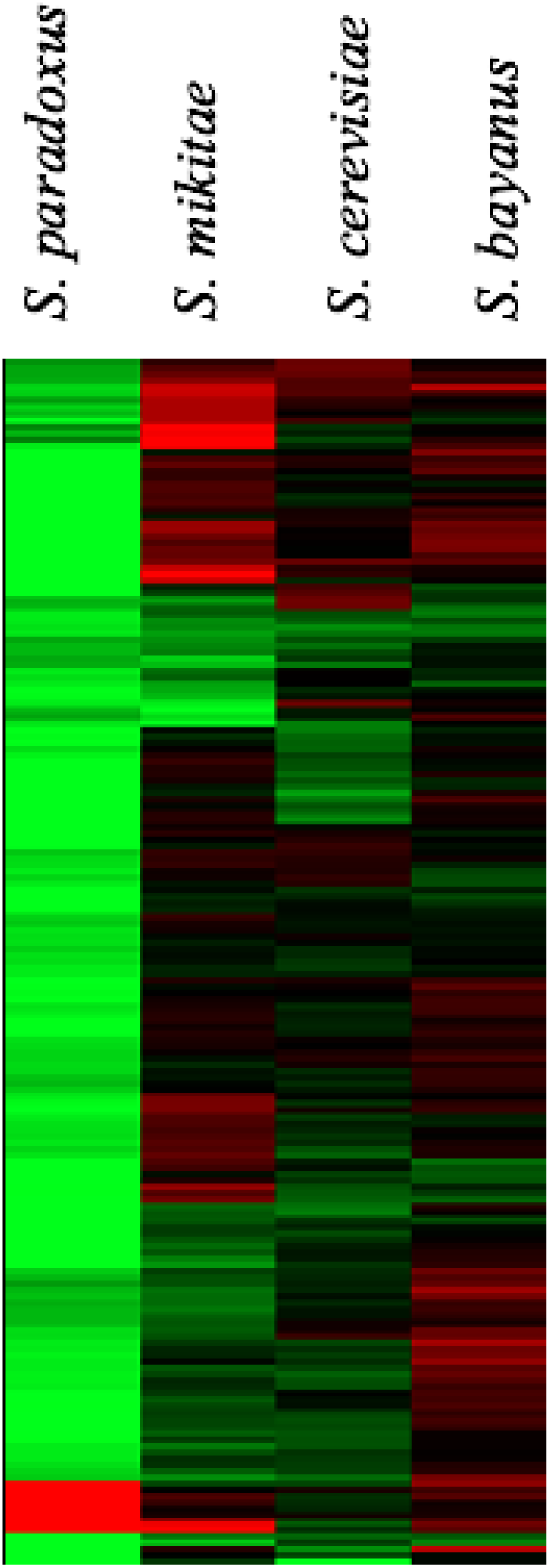
The primary region of repressed genes in *S. paradoxus*. *S. paradoxus* represses genes in this section more broadly and to a greater degree than the other three species.

### Induced and Repressed Gene Overlap

While there is considerable overlap in genes induced between the species, each species has a large number of uniquely induced genes (≥ 1 fold change) (Fig. 5). Furthermore, *S. bayanus* induced far more genes than any of the other species. Overall, there are a greater number of repressed genes than induced between the species in response to MMS (Fig. 5 and Fig. 6). Again, while there is a considerable number of repressed overlapping genes between the species, each species displays uniquely repressed genes (≤ -1 fold change). It is also evident that *S. bayanus* repressed the fewest genes, while *S. paradoxus* repressed the most (Fig. 6).

**Figure 5.**
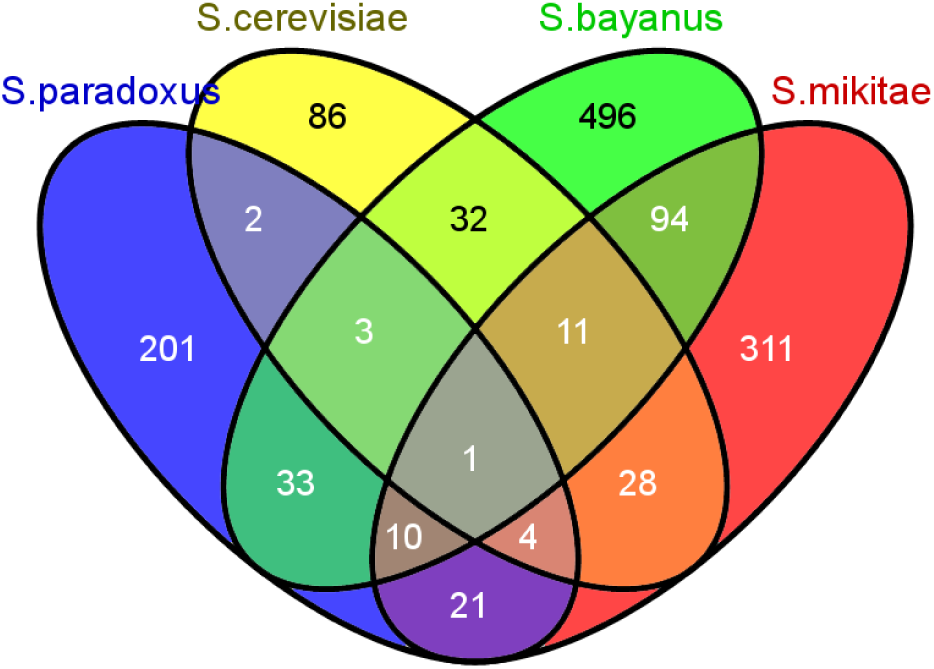
Significantly induced genes (≥ 1 fold change) of the four species in response to MMS.

**Figure 6.**
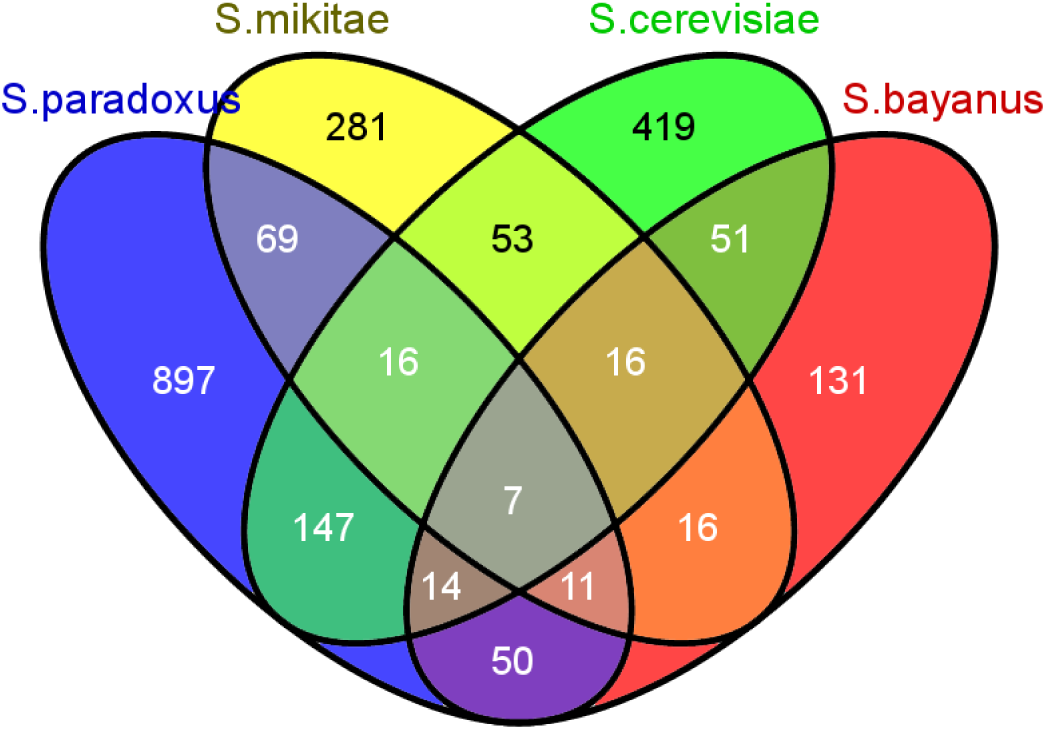
Significantly repressed genes (≤ -1 fold change) of the four species in response to MMS.

### Gene Set Enrichment Analysis

Gene enrichment analysis identified specific processes induced and repressed in response to MMS stress in each species. Gene enrichment allows for an inputted list of genes to be grouped and matched according to particular processes within the cell to identify significant processes (p < 0.05). The results of these analyses are presented in Table 2 to Table 9. *S. bayanus* induced many more processes than the other three species.

### Induced

**Table 2.**
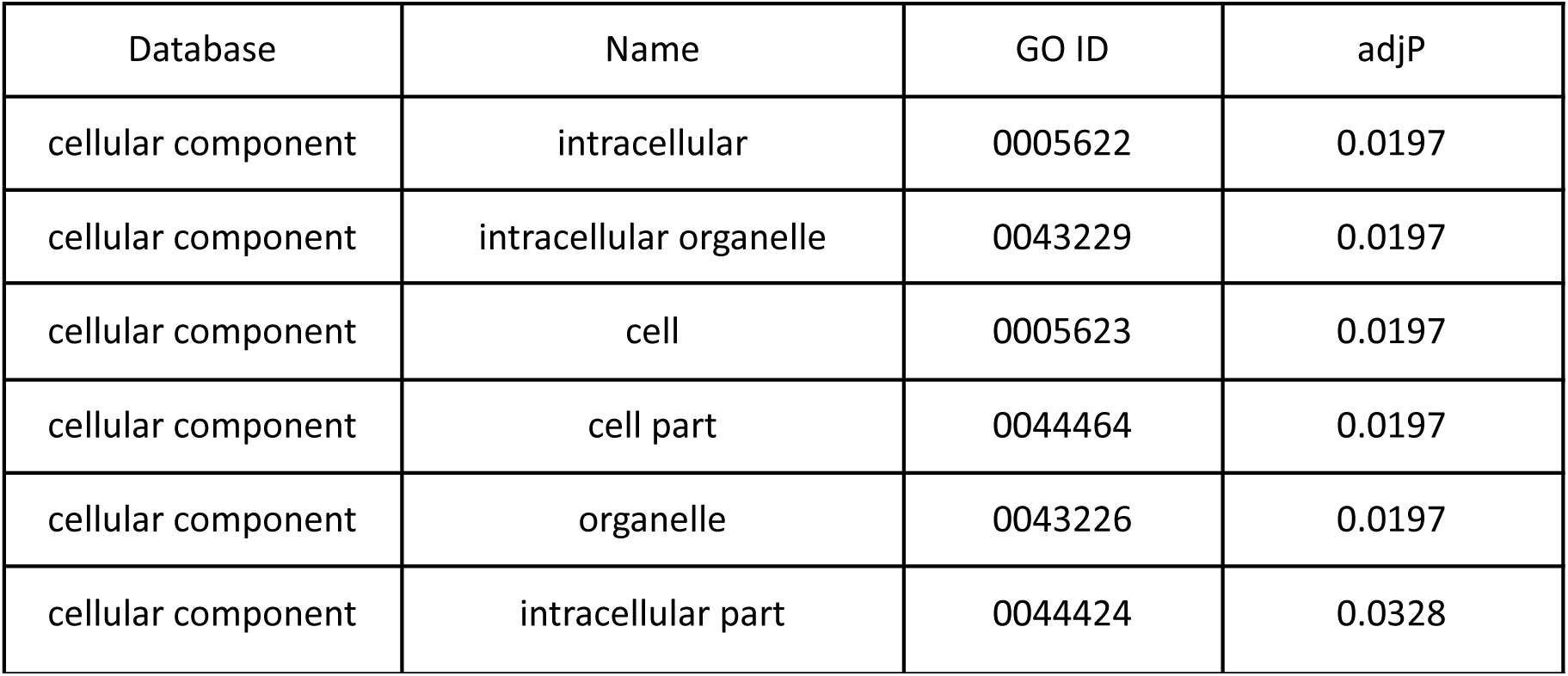
Gene enrichment-identified induced processes in *S. paradoxus* in response to MMS.

| Database | Name | GO ID | adjP |
| --- | --- | --- | --- |
| cellular component | intracellular | 0005622 | 0.0197 |
| cellular component | intracellular organelle | 0043229 | 0.0197 |
| cellular component | cell | 0005623 | 0.0197 |
| cellular component | cell part | 0044464 | 0.0197 |
| cellular component | organelle | 0043226 | 0.0197 |
| cellular component | intracellular part | 0044424 | 0.0328 |

**Table 3.** Gene enrichment-identified induced processes in *S. mikitae* in response to MMS.

| Database | Name | GO ID | adjP |
| --- | --- | --- | --- |
| cellular component | membrane-bounded organelle | 0043227 | 5.09e-09 |
| cellular component | intracellular membrane-bounded organelle | 0043231 | 5.09e-09 |
| cellular component | intracellular organelle | 0043229 | 8.06e-06 |
| cellular component | organelle | 0043226 | 8.06e-06 |
| cellular component | intracellular | 0005622 | 0.0018 |
| cellular component | intracellular part | 0044424 | 0.0025 |
| cellular component | peroxisome | 0005777 | 0.0342 |
| cellular component | nuclear chromosome part | 0044454 | 0.0342 |
| cellular component | nuclear mitotic cohesin complex | 0034990 | 0.0342 |
| cellular component | organelle membrane | 0031090 | 0.0342 |
| cellular component | nucleus | 0005634 | 0.0342 |
| cellular component | chromosomal part | 0044427 | 0.0342 |
| cellular component | microbody | 0042579 | 0.0342 |
| cellular component | mitotic cohesin complex | 0030892 | 0.0342 |
| cellular component | nuclear chromosome | 0000228 | 0.0420 |
| cellular component | chromosome | 0005694 | 0.0425 |

**Table 4.** Gene enrichment-identified induced processes in *S. cerevisiae* in response to MMS.

| Database | Name | GO ID | adjP |
| --- | --- | --- | --- |
| molecular function | oxidoreductase activity | 0016491 | 0.0040 |
| molecular function | N-acetylglucosaminyldiphosphodolichol<br>N-acetylglucosaminyltransferase activity | 0004577 | 0.0476 |

**Table 5.** Gene enrichment-identified induced processes in *S. bayanus* in response to MMS.

| Database | Name | GO ID | adjP |
| --- | --- | --- | --- |
| biological process | protein catabolic process | 0030163 | 7.02e-10 |
| biological process | proteolysis involved in cellular protein catabolic process | 0051603 | 7.52e-10 |
| biological process | modification-dependent protein catabolic process | 0019941 | 8.66e-10 |
| biological process | ubiquitin-dependent protein catabolic process | 0006511 | 8.66e-10 |
| biological process | cellular protein catabolic process | 0044257 | 1.71e-09 |
| biological process | macromolecule catabolic process | 0009057 | 2.06e-08 |
| biological process | modification-dependent macromolecule catabolic process | 0043632 | 2.71e-08 |
| biological process | cellular macromolecule catabolic process | 0044265 | 6.00e-07 |
| biological process | proteasome assembly | 0043248 | 1.91e-06 |
| biological process | proteolysis | 0006508 | 3.67e-06 |
| biological process | proteasomal ubiquitin-dependent protein catabolic process | 0043161 | 6.79e-06 |
| biological process | proteasomal protein catabolic process | 0010498 | 9.88e-06 |
| biological process | cellular response to stress | 0033554 | 0.0002 |
| biological process | response to stress | 0006950 | 0.0002 |
| biological process | mRNA processing | 0006397 | 0.0002 |
| biological process | regulation of protein catabolic process | 0042176 | 0.0002 |
| biological process | protein modification by small protein conjugation or removal | 0070647 | 0.0003 |
| biological process | mRNA splicing, via spliceosome | 0000398 | 0.0004 |
| biological process | cellular response to stimulus | 0051716 | 0.0004 |
| biological process | RNA splicing, via transesterification reactions with bulged adenosine as nucleophile | 0000377 | 0.0004 |
| biological process | response to stimulus | 0050896 | 0.0005 |
| biological process | proteasomal<br>ubiquitin-independent protein<br>catabolic process | 0010499 | 0.0017 |
| biological process | RNA splicing, via transesterification<br>reactions | 0000375 | 0.0018 |
| biological process | protein modification by small<br>protein removal | 0070646 | 0.0020 |
| biological process | RNA splicing | 0008380 | 0.0049 |
| biological process | mRNA metabolic process | 0016071 | 0.0049 |
| biological process | response to oxidative stress | 0006979 | 0.0226 |
| biological process | DNA repair | 0006281 | 0.0226 |
| biological process | flavin-containing compound<br>biosynthetic process | 0042727 | 0.0245 |
| biological process | flavin-containing compound<br>metabolic process | 0042726 | 0.0245 |
| biological process | positive regulation of protein<br>catabolic process | 0045732 | 0.0245 |
| biological process | cellular catabolic process | 0044248 | 0.0260 |
| biological process | iron-sulfur cluster assembly | 0016226 | 0.0260 |
| biological process | metallo-sulfur cluster assembly | 0031163 | 0.0260 |
| biological process | organic substance catabolic process | 1901575 | 0.0289 |
| biological process | catabolic process | 0009056 | 0.0307 |
| biological process | vacuolar transport | 0007034 | 0.0307 |
| biological process | response to DNA damage stimulus | 0006974 | 0.0333 |
| biological process | negative regulation of cellular<br>carbohydrate metabolic process | 0010677 | 0.0379 |
| biological process | negative regulation of carbohydrate metabolic process | 0045912 | 0.0379 |
| molecular function | antioxidant activity | 0016209 | 0.0008 |
| molecular function | threonine-type endopeptidase activity | 0004298 | 0.0040 |
| molecular function | threonine-type peptidase activity | 0070003 | 0.0040 |
| molecular function | peroxidase activity | 0004601 | 0.0164 |
| molecular function | oxidoreductase activity, acting on peroxide as acceptor | 0016684 | 0.0164 |
| molecular function | damaged DNA binding | 0003684 | 0.0273 |
| cellular component | proteasome complex | 0000502 | 1.21e-13 |
| cellular component | proteasome accessory complex | 0022624 | 3.56e-09 |
| cellular component | proteasome regulatory particle | 0005838 | 3.56e-09 |
| cellular component | nucleus | 0005634 | 1.43e-07 |
| cellular component | cytosolic proteasome complex | 0031597 | 3.49e-07 |
| cellular component | proteasome storage granule | 0034515 | 3.49e-07 |
| cellular component | membrane-bounded organelle | 0043227 | 3.71e-06 |
| cellular component | intracellular membrane-bounded organelle | 0043231 | 3.71e-06 |
| cellular component | spliceosomal complex | 0005681 | 1.31e-05 |
| cellular component | U2-type spliceosomal complex | 0005684 | 2.53e-05 |
| cellular component | proteasome regulatory particle, base subcomplex | 0008540 | 6.18e-05 |
| cellular component | proteasome regulatory particle, lid subcomplex | 0008541 | 0.0003 |
| cellular component | prespliceosome | 0071010 | 0.0003 |
| cellular component | U2-type prespliceosome | 0071004 | 0.0003 |
| cellular component | intracellular organelle | 0043229 | 0.0005 |
| cellular component | organelle | 0043226 | 0.0005 |
| cellular component | intracellular | 0005622 | 0.0005 |
| cellular component | intracellular part | 0044424 | 0.0006 |
| cellular component | small nuclear ribonucleoprotein complex | 0030532 | 0.0008 |
| cellular component | proteasome core complex | 0005839 | 0.0009 |
| cellular component | late endosome | 0005770 | 0.0030 |
| cellular component | proteasome core complex, alpha-subunit complex | 0019773 | 0.0043 |
| cellular component | cell part | 0044464 | 0.0175 |
| cellular component | cell | 0005623 | 0.0175 |
| cellular component | protein complex | 0043234 | 0.0175 |
| cellular component | nucleoplasm | 0005654 | 0.0324 |
| cellular component | nuclear part | 0044428 | 0.0370 |
| cellular component | signalosome | 0008180 | 0.0401 |
| cellular component | ribonuclease MRP complex | 0000172 | 0.0409 |
| cellular component | endosome | 0005768 | 0.0437 |

### Repressed

**Table 6.** Gene enrichment-identified repressed processes in *S. paradoxus* in response to MMS.

| Database | Name | GO ID | adjP |
| --- | --- | --- | --- |
| biological process | cellular component organization or biogenesis at cellular level | 0071841 | 6.95e-08 |
| biological process | cellular process | 0009987 | 6.69e-07 |
| biological process | cellular component organization or biogenesis | 0071840 | 2.47e-06 |
| biological process | cellular component organization at cellular level | 0071842 | 2.15e-05 |
| biological process | organelle organization | 0006996 | 3.42e-05 |
| biological process | cellular component organization | 0016043 | 0.0002 |
| biological process | nitrogen compound metabolic process | 0006807 | 0.0041 |
| biological process | mitochondrion organization | 0007005 | 0.0057 |
| biological process | nucleic acid metabolic process | 0090304 | 0.0062 |
| biological process | chromosome segregation | 0007059 | 0.0114 |
| biological process | organic cyclic compound metabolic process | 1901360 | 0.0138 |
| biological process | cellular macromolecular complex subunit organization | 0034621 | 0.0144 |
| biological process | RNA metabolic process | 0016070 | 0.0144 |
| biological process | cellular protein complex assembly | 0043623 | 0.0220 |
| biological process | nucleobase-containing compound metabolic process | 0006139 | 0.0220 |
| biological process | cellular component biogenesis | 0044085 | 0.0220 |
| biological process | cellular aromatic compound metabolic process | 0006725 | 0.0220 |
| biological process | cellular nitrogen compound metabolic process | 0034641 | 0.0280 |
| biological process | heterocycle metabolic process | 0046483 | 0.0280 |
| biological process | amine transport | 0015837 | 0.0280 |
| biological process | macromolecular complex subunit organization | 0043933 | 0.0356 |
| biological process | cellular macromolecular complex assembly | 0034622 | 0.0425 |
| cellular component | intracellular organelle | 0043229 | 1.56e-11 |
| cellular component | organelle | 0043226 | 1.56e-11 |
| cellular component | intracellular membrane-bounded<br>organelle | 0043231 | 2.35e-11 |
| cellular component | membrane-bounded organelle | 0043227 | 2.35e-11 |
| cellular component | intracellular part | 0044424 | 1.04e-10 |
| cellular component | intracellular | 0005622 | 1.04e-10 |
| cellular component | cell part | 0044464 | 1.98e-10 |
| cellular component | cell | 0005623 | 1.98e-10 |
| cellular component | organelle part | 0044422 | 5.15e-08 |
| cellular component | intracellular organelle part | 0044446 | 5.15e-08 |
| cellular component | organelle lumen | 0043233 | 1.63e-06 |
| cellular component | intracellular organelle lumen | 0070013 | 1.63e-06 |
| cellular component | membrane-enclosed lumen | 0031974 | 3.93e-06 |
| cellular component | macromolecular complex | 0032991 | 2.12e-05 |
| cellular component | nucleus | 0005634 | 0.0015 |
| cellular component | nuclear lumen | 0031981 | 0.0028 |
| cellular component | intracellular non-membrane-bounded<br>organelle | 0043232 | 0.0074 |
| cellular component | non-membrane-bounded organelle | 0043228 | 0.0074 |
| cellular component | mitochondrial matrix | 0005759 | 0.0117 |
| cellular component | protein complex | 0043234 | 0.0155 |
| cellular component | nuclear part | 0044428 | 0.0296 |
| cellular component | ribonucleoprotein complex | 0030529 | 0.0343 |
| cellular component | integral to mitochondrial inner membrane | 0031305 | 0.0497 |
| cellular component | organellar small ribosomal subunit | 0000314 | 0.0497 |
| cellular component | mitochondrial small ribosomal subunit | 0005763 | 0.0497 |

**Table 7.** Gene enrichment-identified repressed processes in *S. mikitae* in response to MMS.

| Database | Name | GO ID | adjP |
| --- | --- | --- | --- |
| biological process | organic cyclic compound metabolic process | 1901360 | 3.82e-08 |
| biological process | RNA metabolic process | 0016070 | 3.82e-08 |
| biological process | steroid biosynthetic process | 0006694 | 8.62e-07 |
| biological process | nucleic acid metabolic process | 0090304 | 8.62e-07 |
| biological process | sterol biosynthetic process | 0016126 | 8.62e-07 |
| biological process | sterol metabolic process | 0016125 | 8.84e-07 |
| biological process | steroid metabolic process | 0008202 | 8.84e-07 |
| biological process | ncRNA processing | 0034470 | 1.45e-06 |
| biological process | ncRNA metabolic process | 0034660 | 7.69e-06 |
| biological process | nucleobase-containing compound metabolic process | 0006139 | 7.69e-06 |
| biological process | RNA processing | 0006396 | 7.81e-06 |
| biological process | alcohol metabolic process | 0006066 | 9.21e-06 |
| biological process | cellular aromatic compound metabolic process | 0006725 | 1.13e-05 |
| biological process | heterocycle metabolic process | 0046483 | 1.43e-05 |
| biological process | phytosteroid biosynthetic process | 0016129 | 1.92e-05 |
| biological process | cellular alcohol biosynthetic process | 0044108 | 1.92e-05 |
| biological process | ergosterol biosynthetic process | 0006696 | 1.92e-05 |
| biological process | phytosteroid metabolic process | 0016128 | 4.26e-05 |
| biological process | ergosterol metabolic process | 0008204 | 4.26e-05 |
| biological process | alcohol biosynthetic process | 0046165 | 5.07e-05 |
| biological process | cellular nitrogen compound metabolic process | 0034641 | 7.41e-05 |
| biological process | cellular alcohol metabolic process | 0044107 | 0.0001 |
| biological process | cellular component organization or biogenesis | 0071840 | 0.0001 |
| biological process | organic cyclic compound biosynthetic process | 1901362 | 0.0001 |
| biological process | rRNA processing | 0006364 | 0.0002 |
| biological process | cellular component organization or biogenesis at cellular level | 0071841 | 0.0003 |
| biological process | rRNA metabolic process | 0016072 | 0.0007 |
| biological process | ribosome biogenesis | 0042254 | 0.0007 |
| biological process | RNA biosynthetic process | 0032774 | 0.0008 |
| biological process | transcription, DNA-dependent | 0006351 | 0.0008 |
| biological process | macromolecule methylation | 0043414 | 0.0009 |
| biological process | cytokinetic process | 0032506 | 0.0010 |
| biological process | tRNA methylation | 0030488 | 0.0012 |
| biological process | maturation of 5.8S rRNA from tricistronic rRNA transcript (SSU-rRNA, 5.8S rRNA, LSU-rRNA) | 0000466 | 0.0014 |
| biological process | ribonucleoprotein complex biogenesis | 0022613 | 0.0015 |
| biological process | maturation of 5.8S rRNA | 0000460 | 0.0016 |
| biological process | RNA methylation | 0001510 | 0.0017 |
| biological process | nitrogen compound metabolic process | 0006807 | 0.0017 |
| biological process | transcription from RNA polymerase I promoter | 0006360 | 0.0017 |
| biological process | regulation of transcription, DNA-dependent | 0006355 | 0.0028 |
| molecular function | RNA methyltransferase activity | 0008173 | 0.0025 |
| molecular function | tRNA methyltransferase activity | 0008175 | 0.0025 |
| molecular function | methyltransferase activity | 0008168 | 0.0328 |
| molecular function | snoRNA binding | 0030515 | 0.0328 |
| molecular function | RNA-dependent ATPase activity | 0008186 | 0.0328 |
| molecular function | RNA polymerase II transcription regulatory region sequence-specific DNA binding transcription factor activity involved in negative regulation of transcription | 0001227 | 0.0328 |
| molecular function | RNA polymerase II core promoter proximal region sequence-specific DNA binding transcription factor activity involved in negative regulation of transcription | 0001078 | 0.0328 |
| molecular function | transferase activity, transferring one-carbon groups | 0016741 | 0.0492 |
| molecular function | S-adenosylmethionine-dependent methyltransferase activity | 0008757 | 0.0492 |
| molecular function | transcription factor binding | 0008134 | 0.0492 |
| cellular component | intracellular organelle lumen | 0070013 | 1.07e-07 |
| cellular component | nuclear lumen | 0031981 | 1.07e-07 |
| cellular component | organelle lumen | 0043233 | 1.07e-07 |
| cellular component | membrane-enclosed lumen | 0031974 | 1.07e-07 |
| cellular component | intracellular membrane-bounded organelle | 0043231 | 2.39e-07 |
| cellular component | nucleus | 0005634 | 2.39e-07 |
| cellular component | membrane-bounded organelle | 0043227 | 2.39e-07 |
| cellular component | nucleolus | 0005730 | 3.42e-07 |
| cellular component | organelle | 0043226 | 1.14e-06 |
| cellular component | intracellular organelle | 0043229 | 1.14e-06 |
| cellular component | intracellular part | 0044424 | 4.37e-06 |
| cellular component | nuclear part | 0044428 | 4.37e-06 |
| cellular component | preribosome | 0030684 | 4.99e-06 |
| cellular component | preribosome, large subunit precursor | 0030687 | 4.99e-06 |
| cellular component | intracellular | 0005622 | 8.36e-06 |
| cellular component | cell part | 0044464 | 2.55e-05 |
| cellular component | cell | 0005623 | 2.55e-05 |
| cellular component | non-membrane-bounded organelle | 0043228 | 7.23e-05 |
| cellular component | intracellular non-membrane-bounded organelle | 0043232 | 7.23e-05 |
| cellular component | nucleolar part | 0044452 | 0.0002 |
| cellular component | intracellular organelle part | 0044446 | 0.0002 |
| cellular component | organelle part | 0044422 | 0.0002 |
| cellular component | macromolecular complex | 0032991 | 0.0002 |
| cellular component | Pwp2p-containing subcomplex of 90S preribosome | 0034388 | 0.0047 |
| cellular component | small-subunit processome | 0032040 | 0.0057 |
| cellular component | protein complex | 0043234 | 0.0099 |
| cellular component | DNA-directed RNA polymerase complex | 0000428 | 0.0157 |
| cellular component | nuclear chromatin | 0000790 | 0.0157 |
| cellular component | RNA polymerase complex | 0030880 | 0.0157 |
| cellular component | nucleoplasm | 0005654 | 0.0162 |
| cellular component | chromosome | 0005694 | 0.0193 |
| cellular component | nuclear chromosome | 0000228 | 0.0193 |
| cellular component | cellular bud neck | 0005935 | 0.0193 |
| cellular component | 90S preribosome | 0030686 | 0.0193 |
| cellular component | nucleoplasm part | 0044451 | 0.0195 |
| cellular component | chromosomal part | 0044427 | 0.0200 |
| cellular component | DNA-directed RNA polymerase I complex | 0005736 | 0.0200 |
| cellular component | mitochondrial small ribosomal subunit | 0005763 | 0.0207 |
| cellular component | chromatin | 0000785 | 0.0207 |
| cellular component | organellar small ribosomal subunit | 0000314 | 0.0207 |

**Table 8.** Gene enrichment-identified repressed processes in *S. cerevisiae* in response to MMS.

| Database | Name | GO ID | adjP |
| --- | --- | --- | --- |
| biological process | cytoplasmic translation | 0002181 | 8.41e-17 |
| biological process | cellular component biogenesis at cellular level | 0071843 | 1.81e-07 |
| biological process | ribonucleoprotein complex biogenesis | 0022613 | 4.98e-07 |
| biological process | ribosome biogenesis | 0042254 | 4.98e-07 |
| biological process | nucleoside monophosphate metabolic process | 0009123 | 1.50e-06 |
| biological process | purine nucleoside monophosphate metabolic process | 0009126 | 4.31e-06 |
| biological process | nucleoside monophosphate biosynthetic process | 0009124 | 4.56e-06 |
| biological process | ribonucleoside monophosphate metabolic process | 0009161 | 6.14e-06 |
| biological process | nucleoside phosphate biosynthetic process | 1901293 | 1.05e-05 |
| biological process | purine ribonucleoside monophosphate metabolic process | 0009167 | 1.27e-05 |
| biological process | purine nucleoside monophosphate biosynthetic process | 0009127 | 1.27e-05 |
| biological process | ribonucleoside monophosphate biosynthetic process | 0009156 | 1.74e-05 |
| biological process | purine-containing compound biosynthetic process | 0072522 | 1.88e-05 |
| biological process | rRNA processing | 0006364 | 2.60e-05 |
| biological process | nucleotide biosynthetic process | 0009165 | 3.11e-05 |
| biological process | ribonucleotide biosynthetic process | 0009260 | 3.11e-05 |
| biological process | oxidative phosphorylation | 0006119 | 3.11e-05 |
| biological process | purine ribonucleoside monophosphate biosynthetic process | 0009168 | 4.07e-05 |
| biological process | rRNA metabolic process | 0016072 | 4.27e-05 |
| biological process | purine nucleotide biosynthetic process | 0006164 | 7.68e-05 |
| biological process | mitochondrial ATP synthesis coupled electron transport | 0042775 | 9.20e-05 |
| biological process | ATP synthesis coupled electron transport | 0042773 | 9.20e-05 |
| biological process | cellular process | 0009987 | 9.20e-05 |
| biological process | ribose phosphate biosynthetic process | 0046390 | 0.0001 |
| biological process | RNA processing | 0006396 | 0.0001 |
| biological process | nucleobase biosynthetic process | 0046112 | 0.0001 |
| biological process | IMP metabolic process | 0046040 | 0.0002 |
| biological process | ribosome assembly | 0042255 | 0.0002 |
| biological process | respiratory electron transport chain | 0022904 | 0.0002 |
| biological process | one-carbon metabolic process | 0006730 | 0.0002 |
| biological process | IMP biosynthetic process | 0006188 | 0.0002 |
| biological process | 'de novo' IMP biosynthetic process | 0006189 | 0.0003 |
| biological process | ncRNA processing | 0034470 | 0.0003 |
| biological process | purine ribonucleotide biosynthetic process | 0009152 | 0.0004 |
| biological process | rRNA transport | 0051029 | 0.0005 |
| biological process | organophosphate biosynthetic process | 0090407 | 0.0005 |
| biological process | rRNA export from nucleus | 0006407 | 0.0005 |
| biological process | ribosomal small subunit biogenesis | 0042274 | 0.0005 |
| biological process | electron transport chain | 0022900 | 0.0006 |
| biological process | nucleobase metabolic process | 0009112 | 0.0008 |
| molecular function | structural constituent of ribosome | 0003735 | 3.21e-16 |
| molecular function | structural molecule activity | 0005198 | 3.40e-10 |
| molecular function | rRNA binding | 0019843 | 0.0001 |
| molecular function | monooxygenase activity | 0004497 | 0.0052 |
| molecular function | cyclin-dependent protein kinase regulator activity | 0016538 | 0.0052 |
| molecular function | oxidoreductase activity, acting on paired donors, with incorporation or reduction of molecular oxygen, NADH or NADPH as one donor, and incorporation of one atom of oxygen | 0016709 | 0.0067 |
| molecular function | hydrogen ion transmembrane transporter activity | 0015078 | 0.0101 |
| molecular function | protein kinase regulator activity | 0019887 | 0.0101 |
| molecular function | cytochrome-c oxidase activity | 0004129 | 0.0125 |
| molecular function | heme-copper terminal oxidase activity | 0015002 | 0.0125 |
| molecular function | kinase regulator activity | 0019207 | 0.0125 |
| molecular function | oxidoreductase activity, acting on a heme group of donors | 0016675 | 0.0125 |
| molecular function | oxidoreductase activity, acting on a heme group of donors, oxygen as acceptor | 0016676 | 0.0125 |
| molecular function | monovalent inorganic cation transmembrane transporter activity | 0015077 | 0.0289 |
| molecular function | transferase activity, transferring pentosyl groups | 0016763 | 0.0324 |
| molecular function | ubiquinol-cytochrome-c reductase activity | 0008121 | 0.0362 |
| molecular function | methylenetetrahydrofolate dehydrogenase (NADP+) activity | 0004488 | 0.0362 |
| molecular function | oxidoreductase activity, acting on diphenols and related substances as donors, cytochrome as acceptor | 0016681 | 0.0362 |
| molecular function | oxidoreductase activity, acting on diphenols and related substances as donors | 0016679 | 0.0362 |
| molecular function | C-acyltransferase activity | 0016408 | 0.0425 |
| cellular component | cytosolic ribosome | 0022626 | 2.07e-18 |
| cellular component | ribosomal subunit | 0044391 | 3.42e-15 |
| cellular component | ribosome | 0005840 | 6.01e-15 |
| cellular component | cytosolic part | 0044445 | 2.04e-13 |
| cellular component | ribonucleoprotein complex | 0030529 | 2.04e-13 |
| cellular component | intracellular non-membrane-bounded<br>organelle | 0043232 | 8.56e-13 |
| cellular component | non-membrane-bounded organelle | 0043228 | 8.56e-13 |
| cellular component | cytosolic large ribosomal subunit | 0022625 | 5.81e-11 |
| cellular component | large ribosomal subunit | 0015934 | 1.21e-09 |
| cellular component | cytosolic small ribosomal subunit | 0022627 | 1.04e-08 |
| cellular component | intracellular organelle | 0043229 | 1.04e-08 |
| cellular component | organelle | 0043226 | 1.04e-08 |
| cellular component | intracellular organelle part | 0044446 | 4.86e-06 |
| cellular component | small ribosomal subunit | 0015935 | 4.86e-06 |
| cellular component | organelle part | 0044422 | 4.86e-06 |
| cellular component | organelle inner membrane | 0019866 | 1.37e-05 |
| cellular component | mitochondrial respiratory chain | 0005746 | 1.63e-05 |
| cellular component | macromolecular complex | 0032991 | 2.18e-05 |
| cellular component | respiratory chain | 0070469 | 2.51e-05 |
| cellular component | mitochondrial inner membrane | 0005743 | 2.74e-05 |
| cellular component | cell | 0005623 | 0.0001 |
| cellular component | cell part | 0044464 | 0.0001 |
| cellular component | respiratory chain complex IV | 0045277 | 0.0006 |
| cellular component | mitochondrial respiratory chain complex IV | 0005751 | 0.0006 |
| cellular component | nucleolus | 0005730 | 0.0008 |
| cellular component | preribosome | 0030684 | 0.0010 |
| cellular component | mitochondrial membrane | 0031966 | 0.0011 |
| cellular component | mitochondrial envelope | 0005740 | 0.0011 |
| cellular component | intracellular part | 0044424 | 0.0021 |
| cellular component | respiratory chain complex III | 0045275 | 0.0029 |
| cellular component | intracellular | 0005622 | 0.0029 |
| cellular component | mitochondrial respiratory chain complex III | 0005750 | 0.0029 |
| cellular component | nucleosome | 0000786 | 0.0037 |
| cellular component | mitochondrial membrane part | 0044455 | 0.0045 |
| cellular component | envelope | 0031975 | 0.0051 |
| cellular component | organelle envelope | 0031967 | 0.0051 |
| cellular component | nuclear nucleosome | 0000788 | 0.0090 |
| cellular component | cell division site part | 0032155 | 0.0133 |
| cellular component | cellular bud neck contractile ring | 0000142 | 0.0144 |

**Table 9.** Gene enrichment-identified repressed processes in *S. bayanus* in response to MMS.

| Database | Name | GO ID | adjP |
| --- | --- | --- | --- |
| biological process | cell division | 0051301 | 1.42e-05 |
| biological process | nuclear division | 0000280 | 8.02e-05 |
| biological process | mitosis | 0007067 | 8.02e-05 |
| biological process | M phase of mitotic cell cycle | 0000087 | 9.66e-05 |
| biological process | organelle fission | 0048285 | 0.0001 |
| biological process | mitotic cell cycle | 0000278 | 0.0008 |
| biological process | Microtubule cytoskeleton organization | 0000226 | 0.0015 |
| biological process | steroid metabolic process | 0008202 | 0.0015 |
| biological process | sterol metabolic process | 0016125 | 0.0015 |
| biological process | microtubule-based process | 0007017 | 0.0020 |
| biological process | cytokinesis, completion of separation | 0007109 | 0.0069 |
| biological process | ergosterol biosynthetic process | 0006696 | 0.0106 |
| biological process | cellular alcohol biosynthetic process | 0044108 | 0.0106 |
| biological process | spindle pole body separation | 0000073 | 0.0106 |
| biological process | chromosome segregation | 0007059 | 0.0106 |
| biological process | phytosteroid biosynthetic process | 0016129 | 0.0106 |
| biological process | M phase | 0000279 | 0.0106 |
| biological process | sterol biosynthetic process | 0016126 | 0.0129 |
| biological process | aromatic amino acid family catabolic process | 0009074 | 0.0129 |
| biological process | steroid biosynthetic process | 0006694 | 0.0129 |
| biological process | anaphase | 0051322 | 0.0129 |
| biological process | ergosterol metabolic process | 0008204 | 0.0157 |
| biological process | phytosteroid metabolic process | 0016128 | 0.0157 |
| biological process | cell cycle phase | 0022403 | 0.0181 |
| biological process | cell cycle | 0007049 | 0.0181 |
| biological process | microtubule organizing center organization | 0031023 | 0.0194 |
| biological process | spindle organization | 0007051 | 0.0194 |
| biological process | spindle pole body organization | 0051300 | 0.0194 |
| biological process | cell cycle process | 0022402 | 0.0211 |
| biological process | cellular alcohol metabolic process | 0044107 | 0.0211 |
| biological process | mitotic anaphase B | 0000092 | 0.0292 |
| biological process | cytochrome complex assembly | 0017004 | 0.0319 |
| biological process | cytokinetic cell separation | 0000920 | 0.0319 |
| biological process | phosphorylation | 0016310 | 0.0319 |
| biological process | mitotic spindle organization | 0007052 | 0.0367 |
| biological process | cytokinesis | 0000910 | 0.0367 |
| biological process | microtubule polymerization or depolymerization | 0031109 | 0.0367 |
| biological process | generation of precursor metabolites and energy | 0006091 | 0.0405 |
| biological process | mitotic anaphase | 0000090 | 0.0441 |
| biological process | pyridoxal 5'-phosphate salvage | 0009443 | 0.0498 |
| molecular function | structural constituent of cytoskeleton | 0005200 | 0.0324 |
| molecular function | protein kinase regulator activity | 0019887 | 0.0324 |
| molecular function | kinase regulator activity | 0019207 | 0.0324 |
| cellular component | spindle | 0005819 | 0.0008 |
| cellular component | organelle | 0043226 | 0.0008 |
| cellular component | microtubule cytoskeleton | 0015630 | 0.0008 |
| cellular component | intracellular organelle | 0043229 | 0.0008 |
| cellular component | chromosome, centromeric region | 0000775 | 0.0010 |
| cellular component | intracellular membrane-bounded organelle | 0043231 | 0.0011 |
| cellular component | membrane-bounded organelle | 0043227 | 0.0011 |
| cellular component | cytoskeletal part | 0044430 | 0.0013 |
| cellular component | kinetochore | 0000776 | 0.0013 |
| cellular component | cytoskeleton | 0005856 | 0.0021 |
| cellular component | condensed chromosome kinetochore | 0000777 | 0.0021 |
| cellular component | condensed chromosome, centromeric region | 0000779 | 0.0036 |
| cellular component | microtubule | 0005874 | 0.0036 |
| cellular component | microtubule organizing center part | 0044450 | 0.0051 |
| cellular component | inner plaque of spindle pole body | 0005822 | 0.0063 |
| cellular component | spindle pole | 0000922 | 0.0089 |
| cellular component | microtubule organizing center | 0005815 | 0.0131 |
| cellular component | spindle pole body | 0005816 | 0.0131 |
| cellular component | extracellular region | 0005576 | 0.0186 |
| cellular component | organelle part | 0044422 | 0.0275 |
| cellular component | intracellular organelle part | 0044446 | 0.0275 |
| cellular component | cellular bud | 0005933 | 0.0275 |
| cellular component | U2-type spliceosomal complex | 0005684 | 0.0275 |
| cellular component | cellular bud neck | 0005935 | 0.0275 |
| cellular component | cell | 0005623 | 0.0300 |
| cellular component | cell part | 0044464 | 0.0300 |
| cellular component | nucleus | 0005634 | 0.0332 |
| cellular component | respiratory chain | 0070469 | 0.0396 |
| cellular component | gamma-tubulin small complex | 0008275 | 0.0487 |
| cellular component | nuclear chromosome, telomeric region | 0000784 | 0.0487 |
| cellular component | gamma-tubulin small complex, spindle pole body | 0000928 | 0.0487 |
| cellular component | gamma-tubulin complex | 0000930 | 0.0487 |

### Promoter Analysis

Promoter analysis of highly induced (≥ 3 folds) and highly repressed (≤ -3 folds) genes identified several transcription factors. The induced transcription factors have functions that are associated with maintaining the integrity of DNA or affecting the cell cycle. The repressed transcription factors were more varied in their functions.

**Table 10.** Induced transcription factors in response to MMS (Induced MMS Transcription Factors).

| Induced Transcription Factors | Function |
| --- | --- |
| Abf1p | DNA binding protein involved in DNA replication and repair |
| Azf1p | Genetic and physical interactions suggest role in genome maintenance |
| Hme1p | Forkhead transcription factor that drives S-phase specific expression of genes involved in chromosome segregation, spindle dynamics, and budding |
| Rfx1p | Involved in DNA damage and replication checkpoint pathway |
| Swi4p | Regulates late G1-specific transcription of targets including cyclins and genes required for DNA synthesis and repair |

**Table 11.**
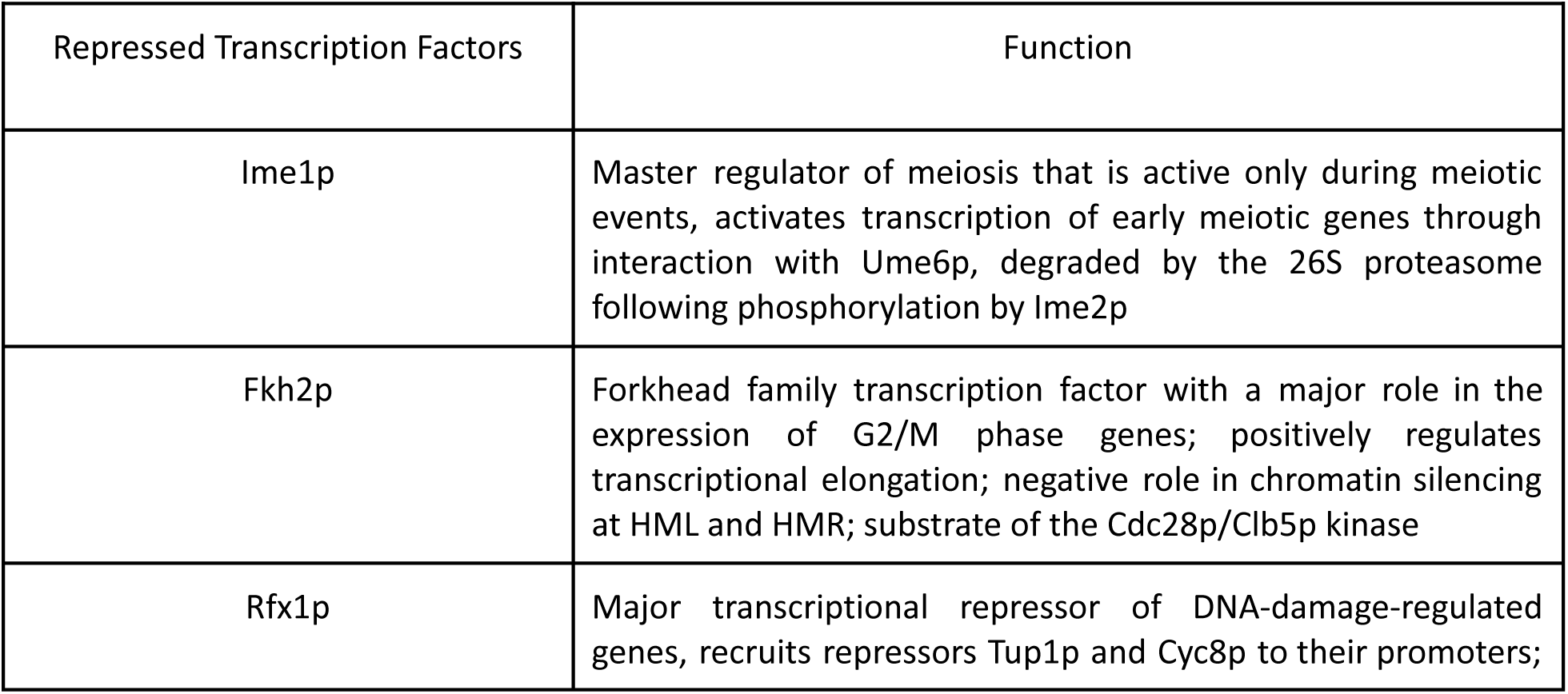
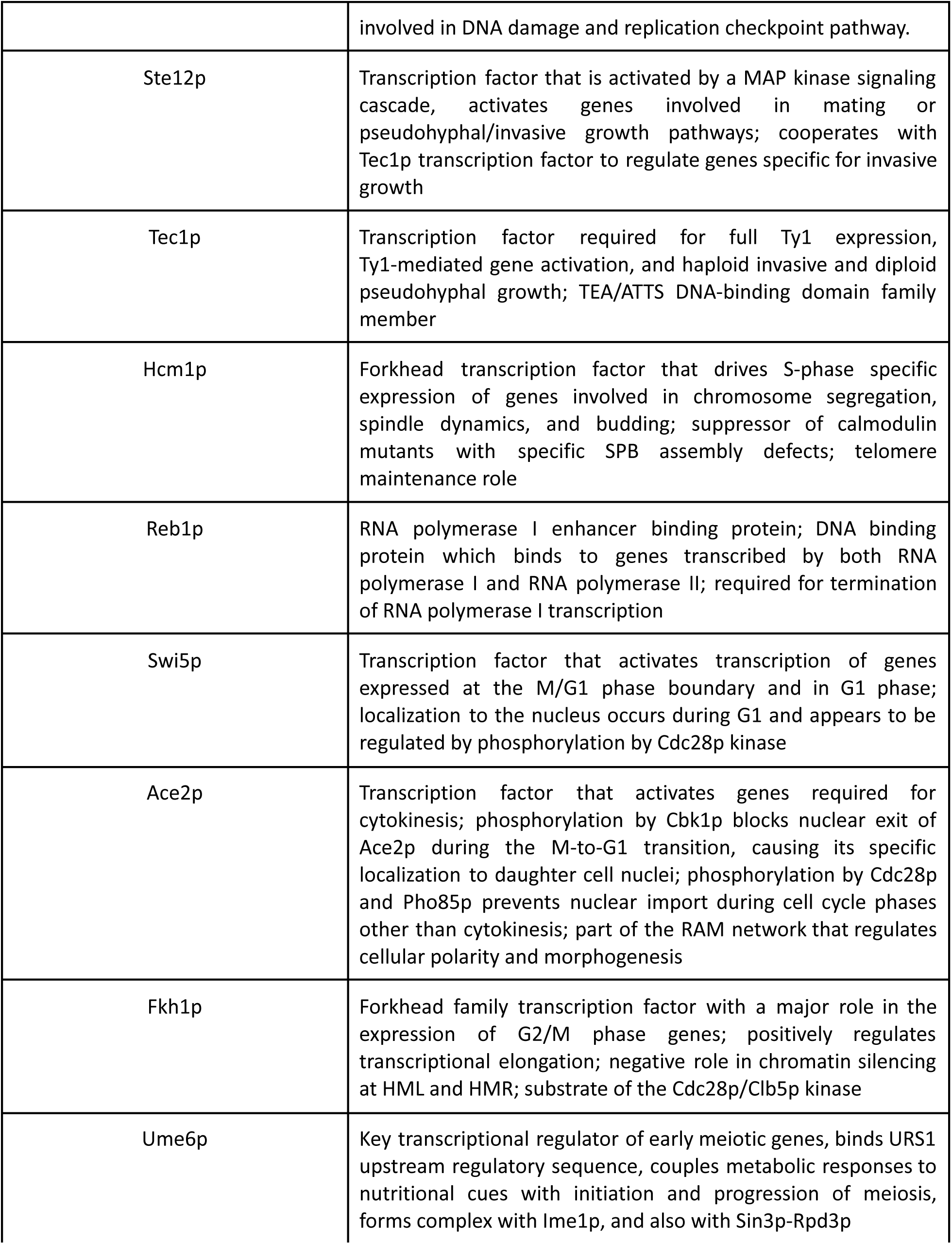
Gene enrichment-identified repressed processes in *S. bayanus* in response to MMS.

### Concentration Gradient Response

The concentration gradient response of *S. cerevisiae* showed that the fold changes in the highly induced genes tested in *S. cerevisiae* do not plateau for genes tested (RAD54, DIN7, and IRC19) for the tested concentrations of MMS (Fig. 7). MMS concentrations above 0.06% did not allow for yeast growth in differential growth characterizations, thus this concentration was utilized as the upper limit for the gradient response experiment. A positive correlations was noted between the concentration of MMS used and the fold change in highly induced genes. Pearson’s R values for RAD54, DIN7, and IRC19 were 0.7995, 0.9539, and 0.9550, respectively.

**Figure 7.**
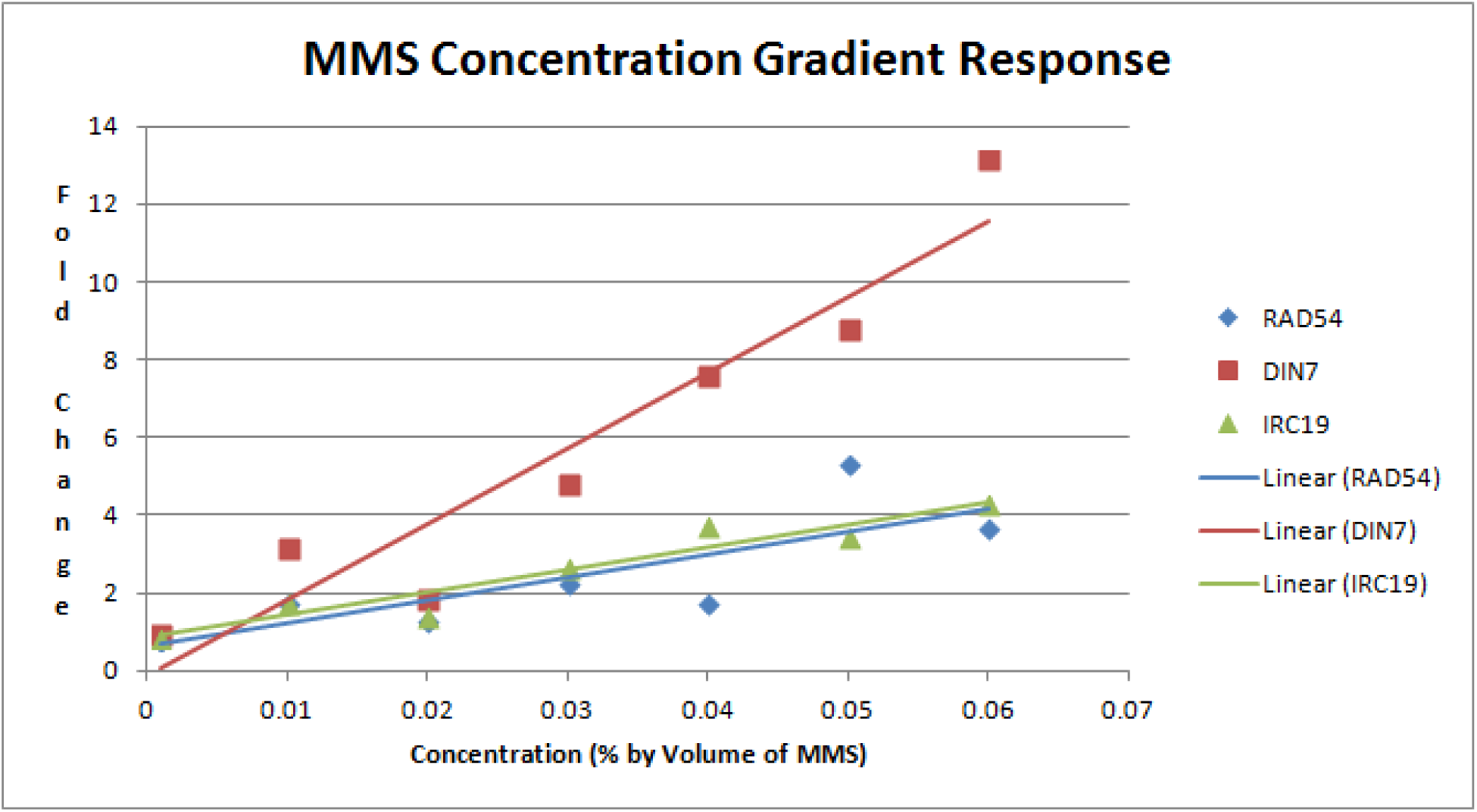
Concentration dependent response of RAD54, DIN7, and IRC16 in *S. cerevisiae*.

## Discussion

The different growth abilities of yeast species in response to a stress condition is association with their capability to vary genomic expression to facilitate repair and maintenance of genomic material. Prior literature shows that MMS-induced DNA damage results in rapid and extensive changes in genomic expression in wild type species (Jelinsky and Samson, 1999). Only cells capable of maintaining genomic integrity through, in part, appropriate genomic expression changes will propagate. In this study, *S. bayanus* did not show any growth in the stress condition. This is particularly interesting since *S. bayanus* is the most divergent species in our study (Scannell et al., 2011). This may indicate an ancestral state that is unable to respond adequately to DNA damage induced by MMS stress.

At the level of gene expression, this study depicted genome-wide expression patterns in response to MMS. Despite considerable overlap in highly induced and repressed genes, the organisms were observed to have vastly differing and unique expression patterns (Fig _ and Fig _). This finding supports the notion that different yeast species have different ways of altering their expression patterns in response to stresses as presented by Tirosh et al (http://www.pnas.org/content/108/40/16693.full.pdf). However, what these changes are and their implications are of interest. *S. paradoxus*, which grew well under the MMS condition, demonstrated sections in its transcriptome that were highly repressed (Fig. 4). Our findings demonstrate that this species not only repressed more genes, but also does so more significantly than the other species. In contrast, *S. bayanus* proportionally had the highest ratio of highly induced genes and demonstrated the least number of highly repressed genes (Fig.2, Fig.5, and Fig. 6). This is in stark contrast to the expression pattern of *S. paradoxus* and may serve to explain the poor growth of *S. bayanus* in the stress condition. Over the course of evolution, the specific ways in which these species respond to DNA-damage have altered. The results of the gene enrichment and gene overlap analyses lead the authors to propose that an appropriate response to DNA-damage from MMS stress induces a small subset of genes, including those essential for DNA repair and maintenance, and represses a far larger number of genes. This may be a mechanism by which the cell conserves resources by repressing many processes and channels its efforts into inducing a smaller number of genes as part of the stress response (Vilaprinyo et al., 2006). This theory has the potential to alter how the results of future stress responses are viewed and conducted. Furthermore, it emphasizes the importance of determining the functional significance of gene expression, such as through gene enrichment analyses.

Transcription factors in *S. cerevisiae* have been studied as part of general stress response pathways (Estruch, 2000). In this study, we obtained a set of induced and repressed transcription factors found to be highly induced/repressed not only in *S. cerevisiae*, but also in the other three species. We named them the MMS Response Transcription Factors. This presents an overarching set of significant MMS response transcription factors for multiple species within the *sensu stricto* genus of *Saccharomyces* yeast. The MMS Response Transcription Factors have the potential to broaden the scope of investigation for subsequent DNA damage experiments in *Saccharomyces* yeast and steer the direction of future studies by identifying more common principles of stress response to MMS.

Yeast respond to mating pheromone concentration gradients by projecting up the gradient (Arkowitz, 2009). Furthermore, yeast have been shown to be alter their response in accordance to the concentration gradient (Moore et al., 2013). It is also known that *S. cerevisiae* induces the expression of specific genes in response to DNA-damaging agents such as MMS in its environment (Ruby and Szostak, 1985). We investigated whether the gene expression response of *S. cerevisiae* would demonstrate dose-dependence to MMS. This analysis demonstrated that *S. cerevisiae* shows strong, positive, and linear correlations between the fold changes of induced genes and increasing concentrations of MMS. A plateau was not observed for the doses tested. This finding lends us to believe that in a stress response it is not only important to uniquely regulate genome expression, but that how greatly regulated the genes are also confers a successful stress response.

There are a few limitations of this study at present. First, as a preliminary study, this investigation requires replication to validate results. Second, the investigation is unable to delineate the effects of specific pathways in the MMS response. As such, it may be bolstered by knock-out experiments of particular genes implicated in the MMS stress response, such as RAD54, in order to observe the resulting gene expression effects. Third, since concentration gradient studies were only performed for *S. cerevisiae* (as it is the model yeast species), the findings may not be applicable to the other species, warranting further investigation.

## Conclusion [to be done]

## Supplemental transcription factor data

### Induced

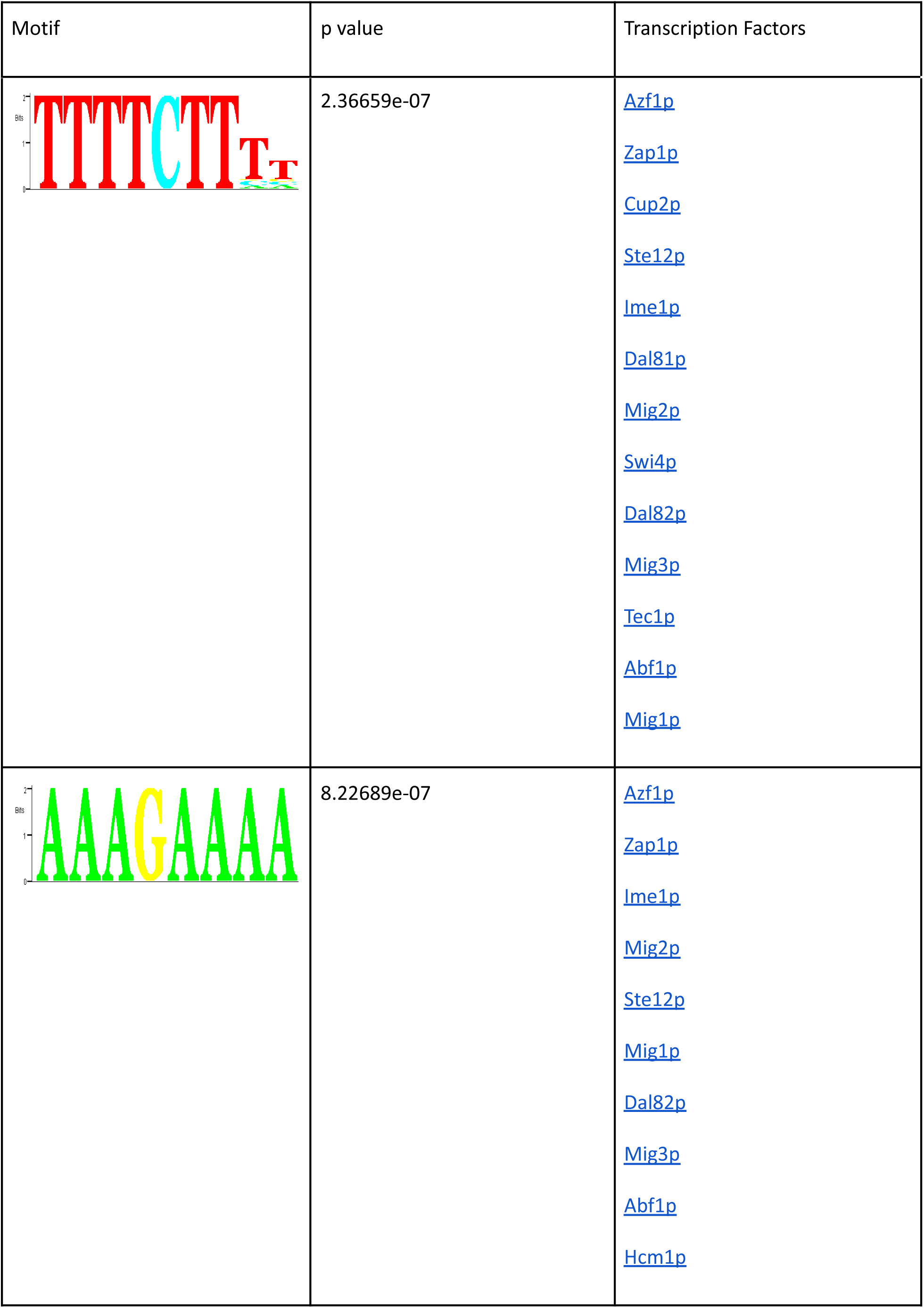
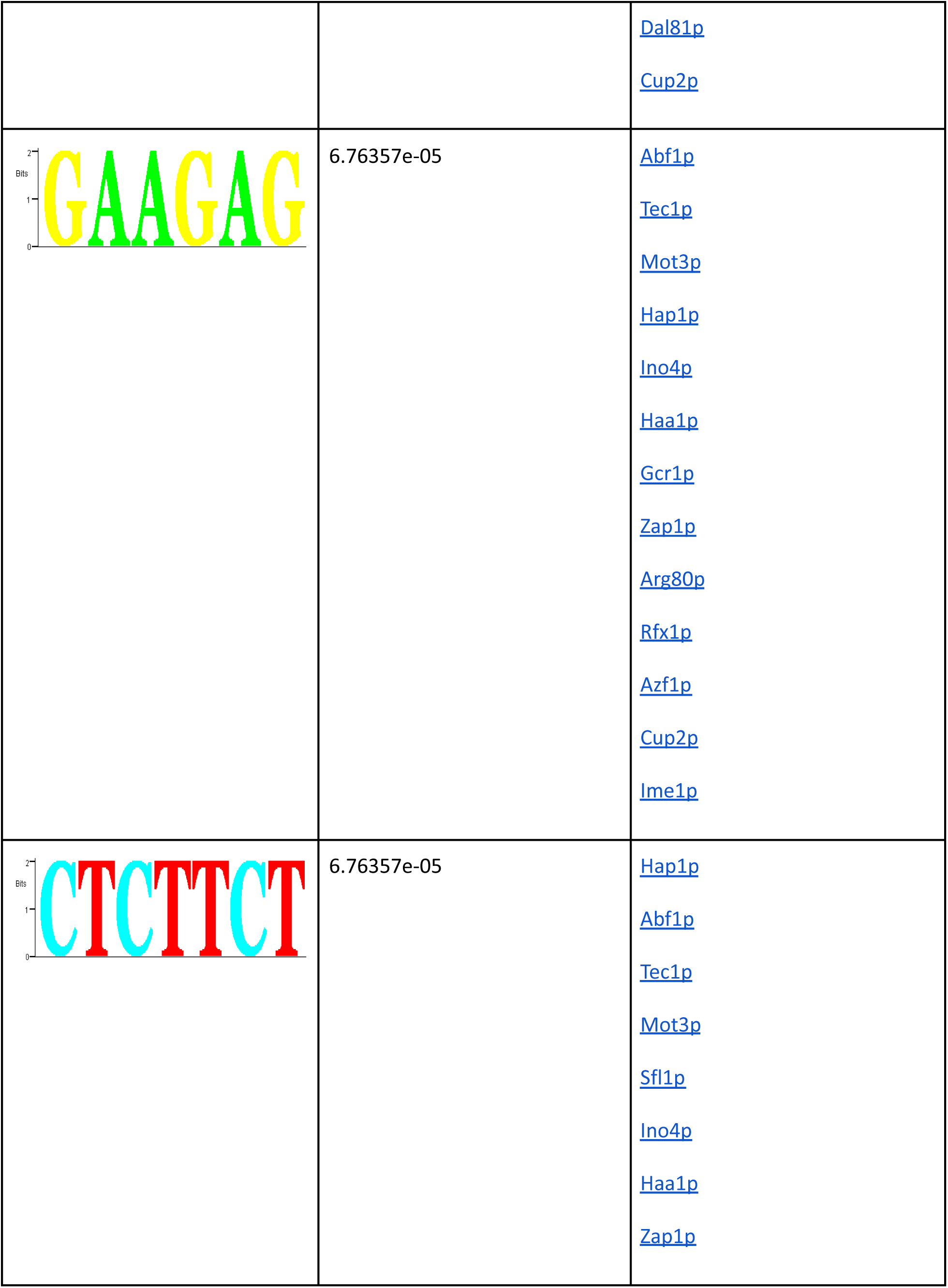
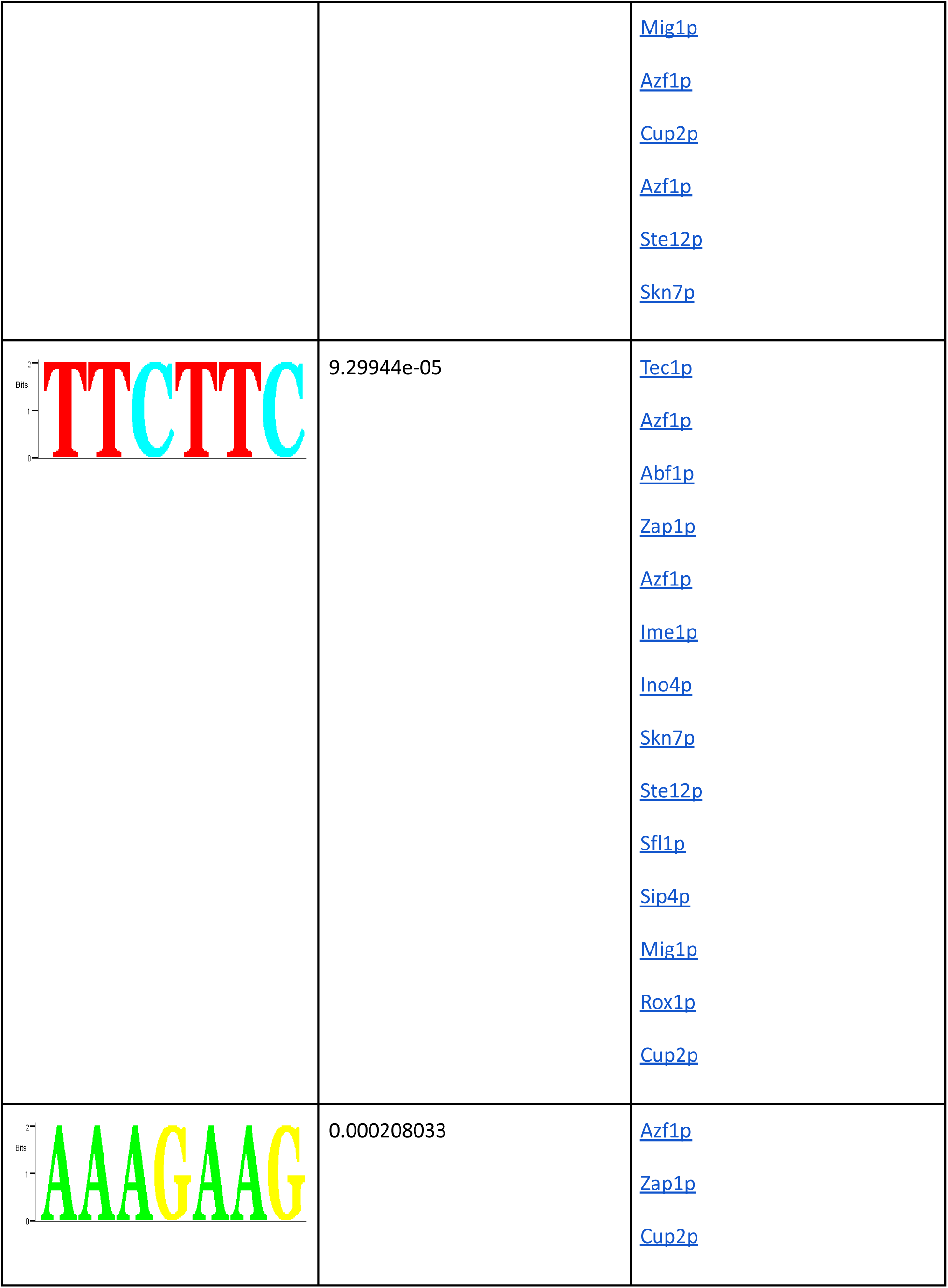
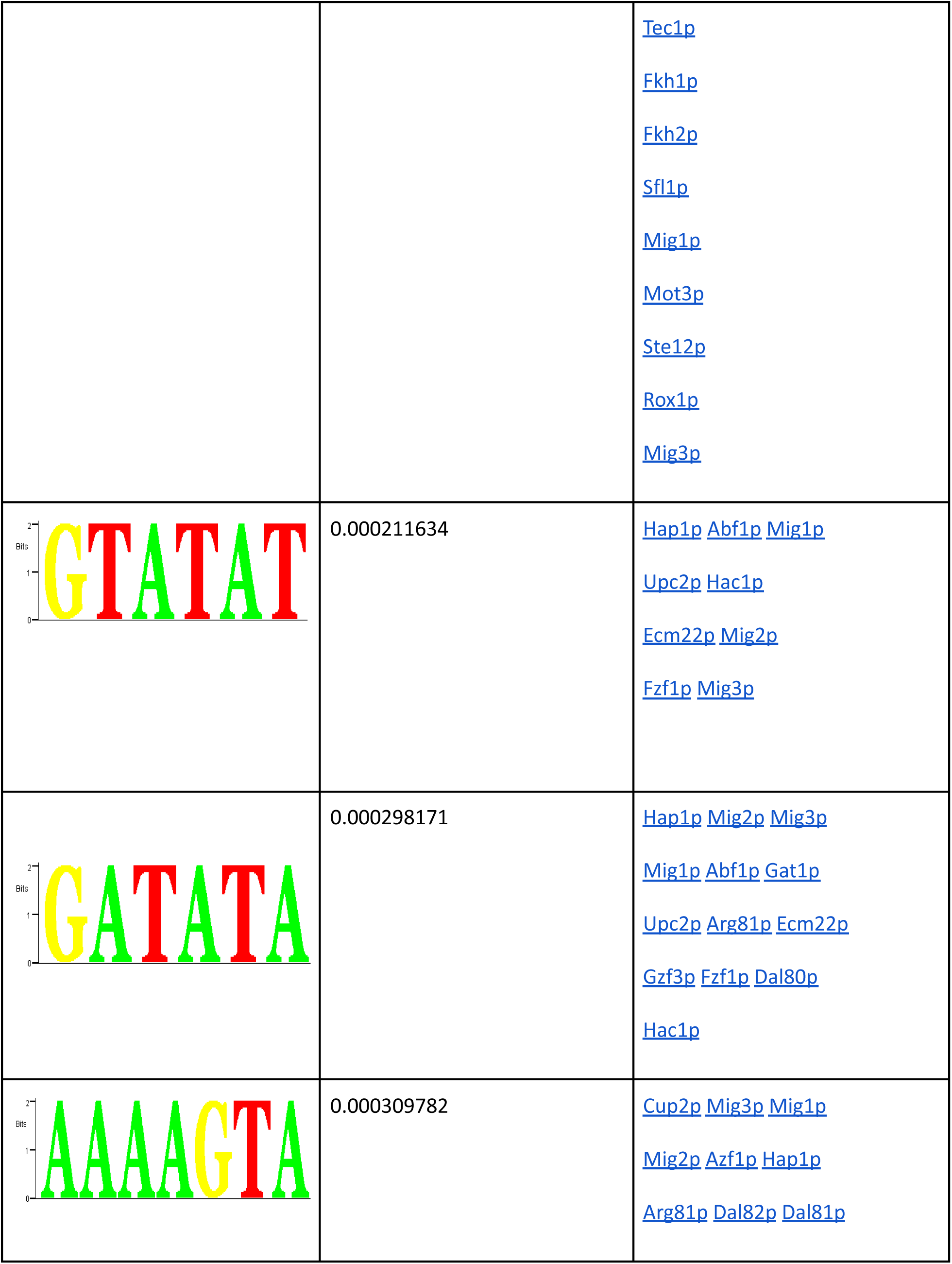
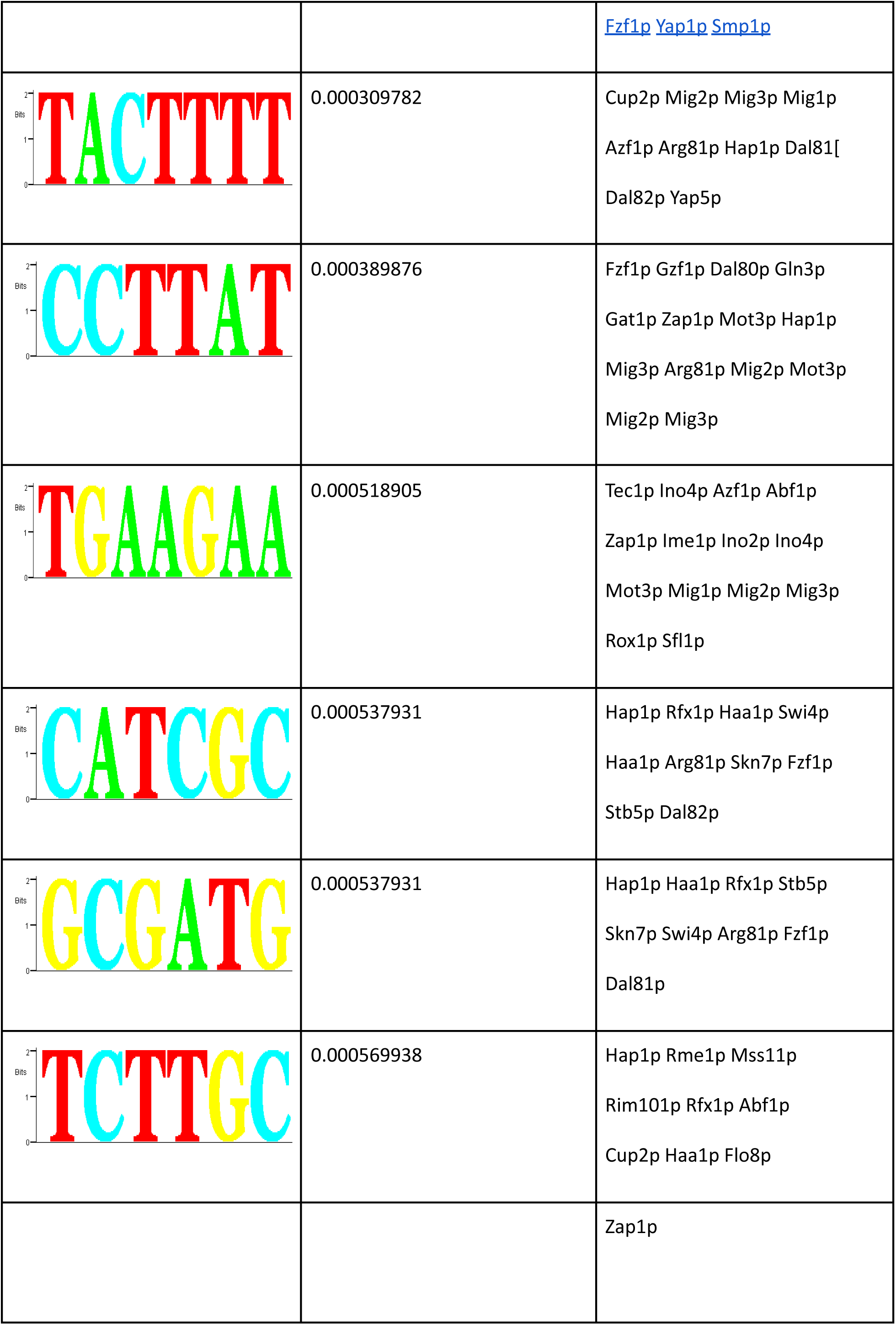

### Repressed

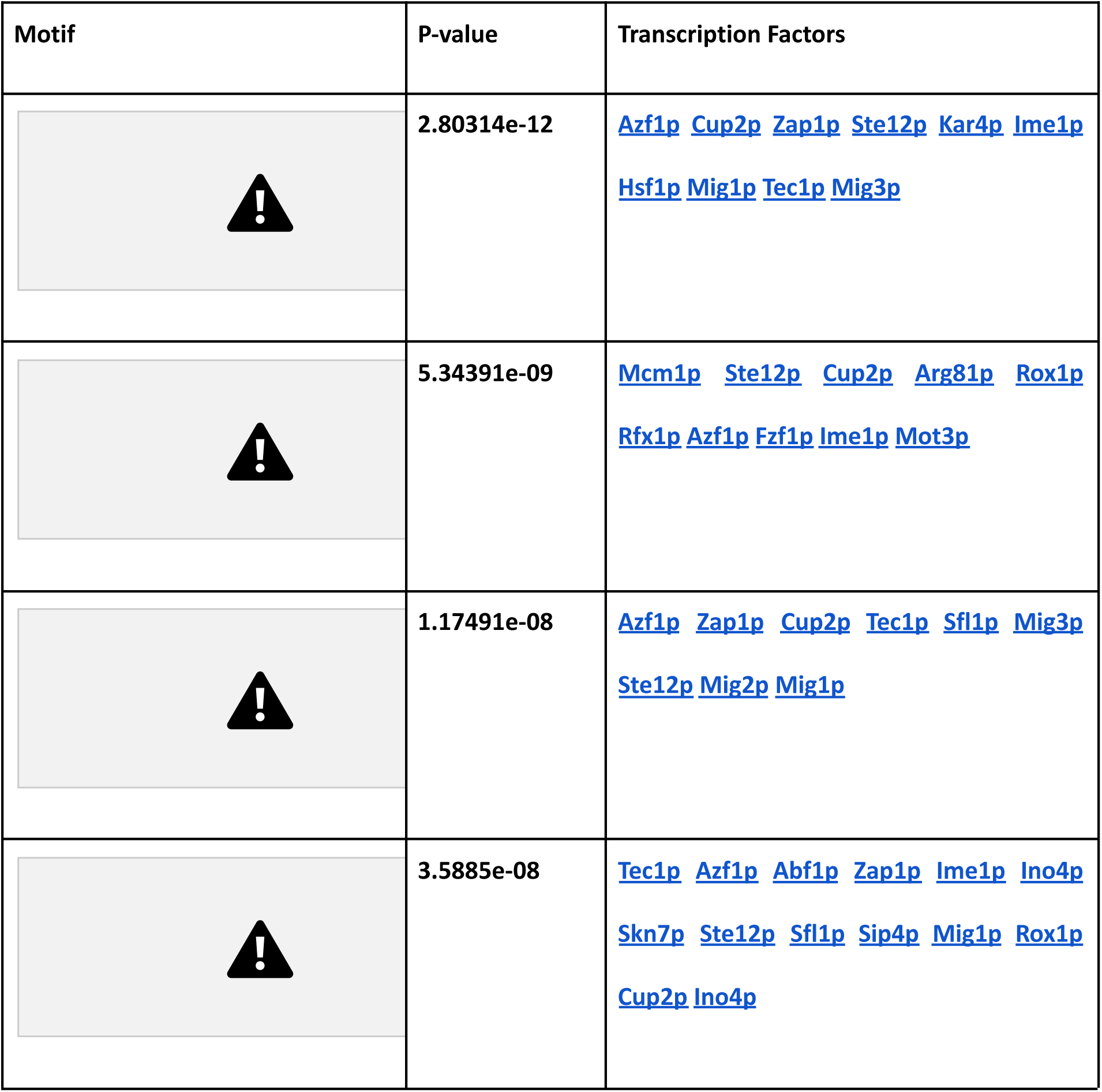
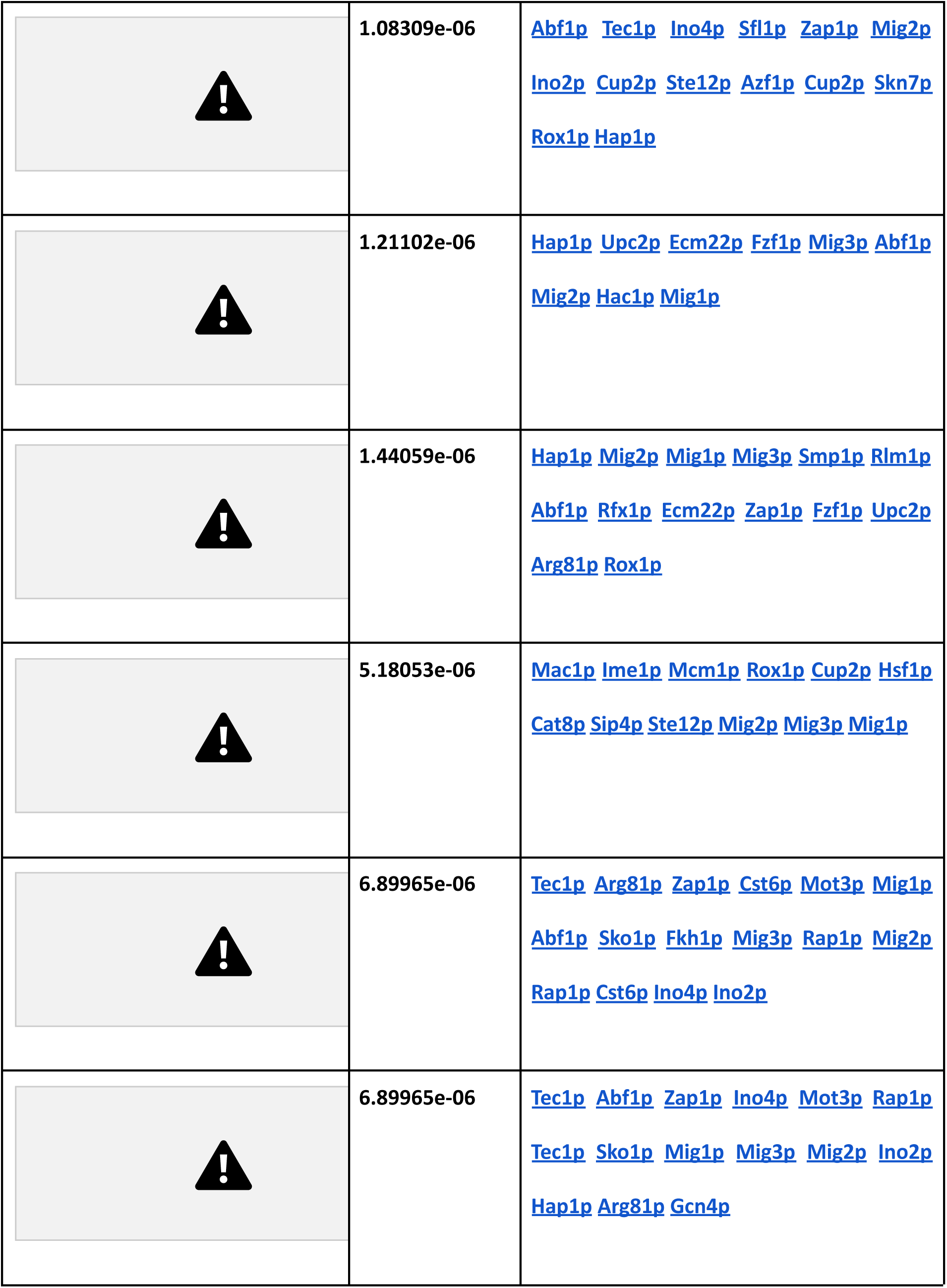
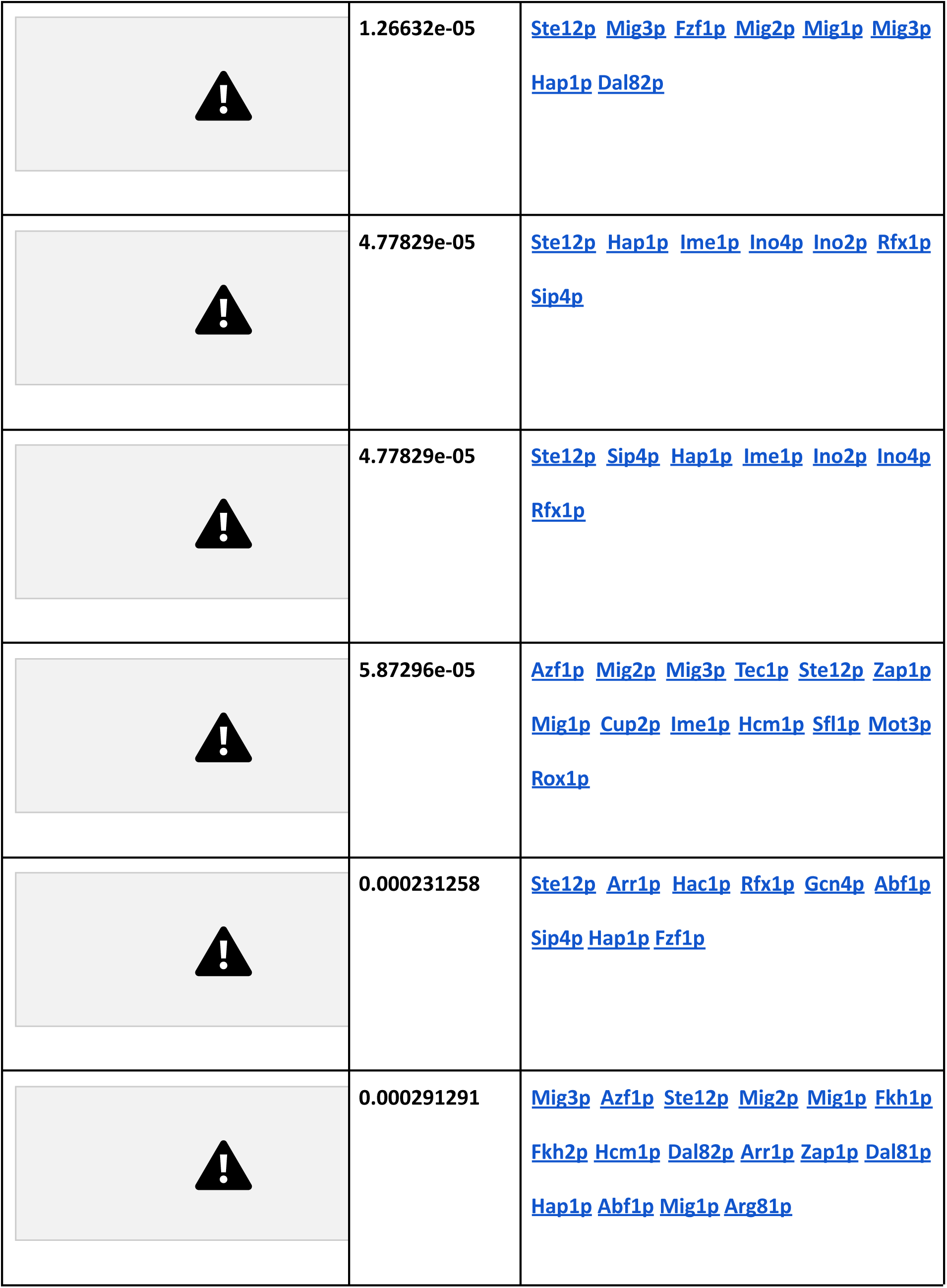
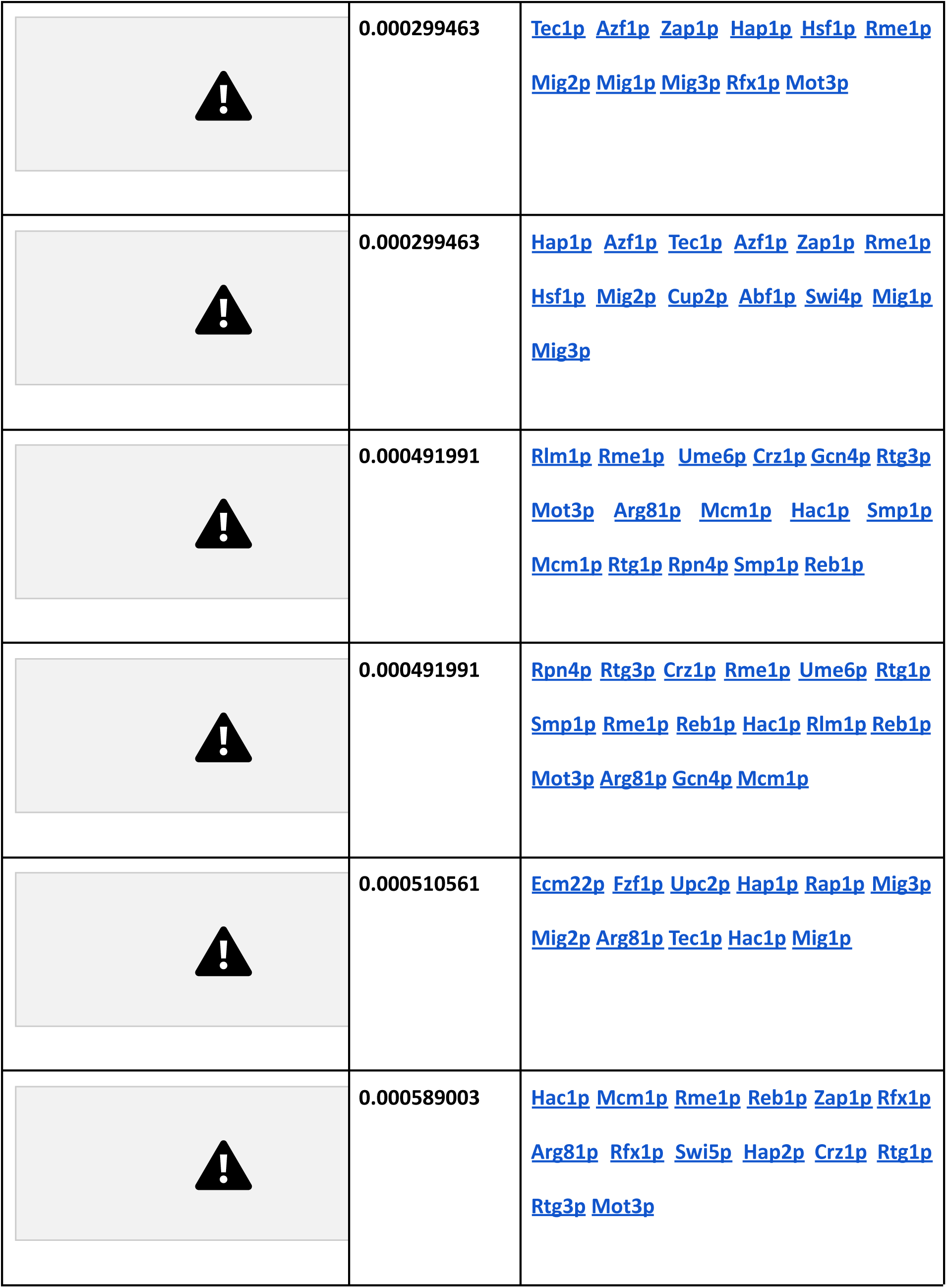
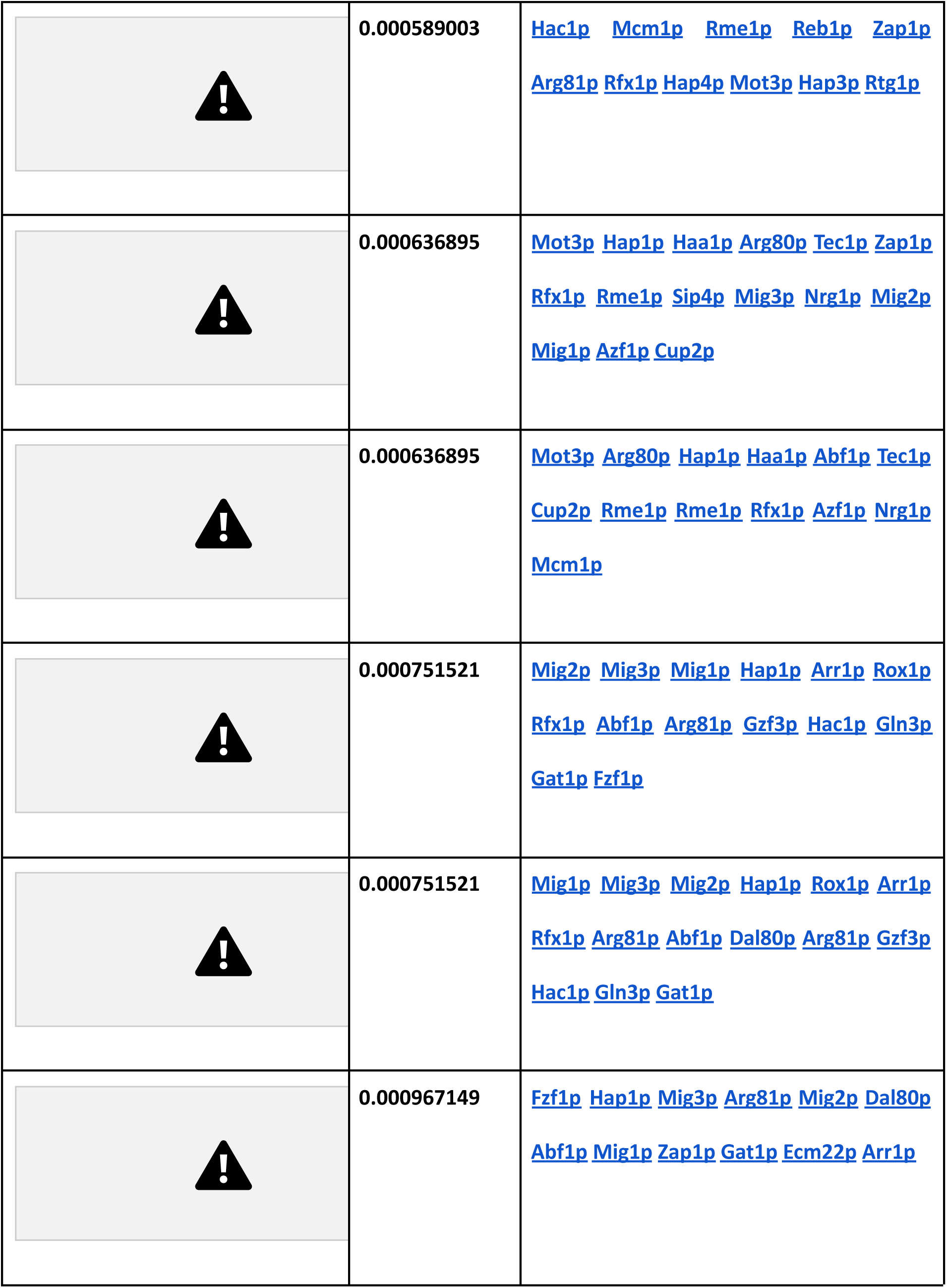

| Motif | P-value | Transcription Factors |
| --- | --- | --- |
|  | 2.80314e-12 | Azf1p Cup2p Zap1p Ste12p Kar4p Ime1p<br>Hsf1p Mig1p Tec1p Mig3p |
|  | 5.34391e-09 | Mcm1p Ste12p Cup2p Arg81p Rox1p<br>Rfx1p Azf1p Fzf1p Ime1p Mot3p |
|  | 1.17491e-08 | Azf1p Zap1p Cup2p Tec1p Sfl1p Mig3p<br>Ste12p Mig2p Mig1p |
|  | 3.5885e-08 | Tec1p Azf1p Abf1p Zap1p Ime1p Ino4p<br>Skn7p Ste12p Sfl1p Sip4p Mig1p Rox1p<br>Cup2p Ino4p |
|  | 1.08309e-06 | Abf1p Tec1p Ino4p Sfl1p Zap1p Mig2p<br>Ino2p Cup2p Ste12p Azf1p Cup2p Skn7p<br>Rox1p Hap1p |
|  | 1.21102e-06 | Hap1p Upc2p Ecm22p Fzf1p Mig3p Abf1p<br>Mig2p Hac1p Mig1p |
|  | 1.44059e-06 | Hap1p Mig2p Mig1p Mig3p Smp1p Rlm1p<br>Abf1p Rfx1p Ecm22p Zap1p Fzf1p Upc2p<br>Arg81p Rox1p |
|  | 5.18053e-06 | Mac1p Ime1p Mcm1p Rox1p Cup2p Hsf1p<br>Cat8p Sip4p Ste12p Mig2p Mig3p Mig1p |
|  | 6.89965e-06 | Tec1p Arg81p Zap1p Cst6p Mot3p Mig1p<br>Abf1p Sko1p Fkh1p Mig3p Rap1p Mig2p<br>Rap1p Cst6p Ino4p Ino2p |
|  | 6.89965e-06 | Tec1p Abf1p Zap1p Ino4p Mot3p Rap1p<br>Tec1p Sko1p Mig1p Mig3p Mig2p Ino2p<br>Hap1p Arg81p Gcn4p |
|  | 1.26632e-05 | Ste12p Mig3p Fzf1p Mig2p Mig1p Mig3p<br>Hap1p Dal82p |
|  | 4.77829e-05 | Ste12p Hap1p Ime1p Ino4p Ino2p Rfx1p<br>Sip4p |
|  | 4.77829e-05 | Ste12p Sip4p Hap1p Ime1p Ino2p Ino4p<br>Rfx1p |
|  | 5.87296e-05 | Azf1p Mig2p Mig3p Tec1p Ste12p Zap1p<br>Mig1p Cup2p Ime1p Hcm1p Sfl1p Mot3p<br>Rox1p |
|  | 0.000231258 | Ste12p Arr1p Hac1p Rfx1p Gcn4p Abf1p<br>Sip4p Hap1p Fzf1p |
|  | 0.000291291 | Mig3p Azf1p Ste12p Mig2p Mig1p Fkh1p<br>Fkh2p Hcm1p Dal82p Arr1p Zap1p Dal81p<br>Hap1p Abf1p Mig1p Arg81p |
|  | 0.000299463 | Tec1p Azf1p Zap1p Hap1p Hsf1p Rme1p<br>Mig2p Mig1p Mig3p Rfx1p Mot3p |
|  | 0.000299463 | Hap1p Azf1p Tec1p Azf1p Zap1p Rme1p<br>Hsf1p Mig2p Cup2p Abf1p Swi4p Mig1p<br>Mig3p |
|  | 0.000491991 | Rlm1p Rme1p Ume6p Crz1p Gcn4p Rtg3p<br>Mot3p Arg81p Mcm1p Hac1p Smp1p<br>Mcm1p Rtg1p Rpn4p Smp1p Reb1p |
|  | 0.000491991 | Rpn4p Rtg3p Crz1p Rme1p Ume6p Rtg1p<br>Smp1p Rme1p Reb1p Hac1p Rlm1p Reb1p<br>Mot3p Arg81p Gcn4p Mcm1p |
|  | 0.000510561 | Ecm22p Fzf1p Upc2p Hap1p Rap1p Mig3p<br>Mig2p Arg81p Tec1p Hac1p Mig1p |
|  | 0.000589003 | Hac1p Mcm1p Rme1p Reb1p Zap1p Rfx1p<br>Arg81p Rfx1p Swi5p Hap2p Crz1p Rtg1p<br>Rtg3p Mot3p |
|  | 0.000589003 | Hac1p Mcm1p Rme1p Reb1p Zap1p<br>Arg81p Rfx1p Hap4p Mot3p Hap3p Rtg1p |
|  | 0.000636895 | Mot3p Hap1p Haa1p Arg80p Tec1p Zap1p<br>Rfx1p Rme1p Sip4p Mig3p Nrg1p Mig2p<br>Mig1p Azf1p Cup2p |
|  | 0.000636895 | Mot3p Arg80p Hap1p Haa1p Abf1p Tec1p<br>Cup2p Rme1p Rme1p Rfx1p Azf1p Nrg1p<br>Mcm1p |
|  | 0.000751521 | Mig2p Mig3p Mig1p Hap1p Arr1p Rox1p<br>Rfx1p Abf1p Arg81p Gzf3p Hac1p Gln3p<br>Gat1p Fzf1p |
|  | 0.000751521 | Mig1p Mig3p Mig2p Hap1p Rox1p Arr1p<br>Rfx1p Arg81p Abf1p Dal80p Arg81p Gzf3p<br>Hac1p Gln3p Gat1p |
|  | 0.000967149 | Fzf1p Hap1p Mig3p Arg81p Mig2p Dal80p<br>Abf1p Mig1p Zap1p Gat1p Ecm22p Arr1p |

### Repressed Transcription Factors

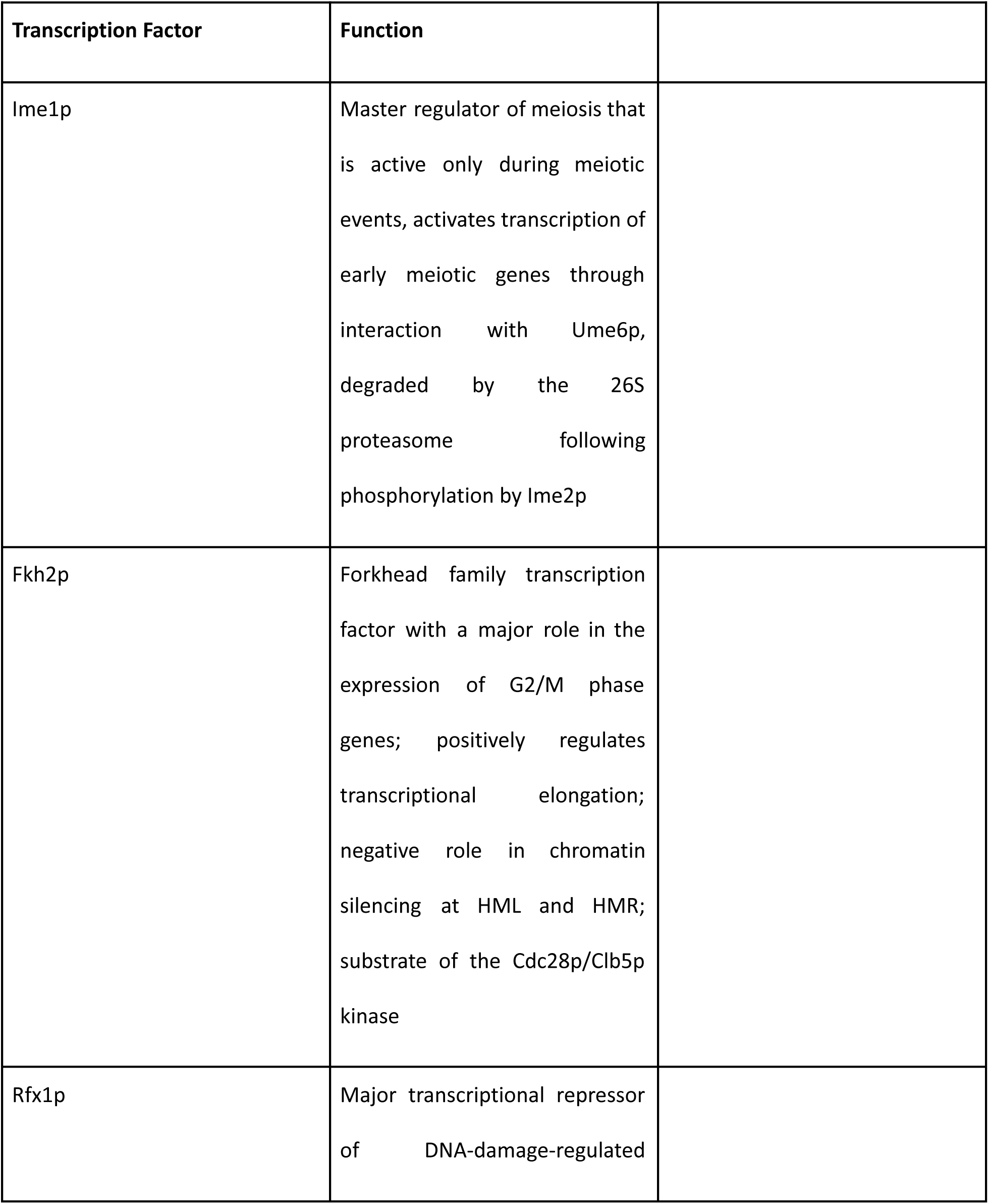
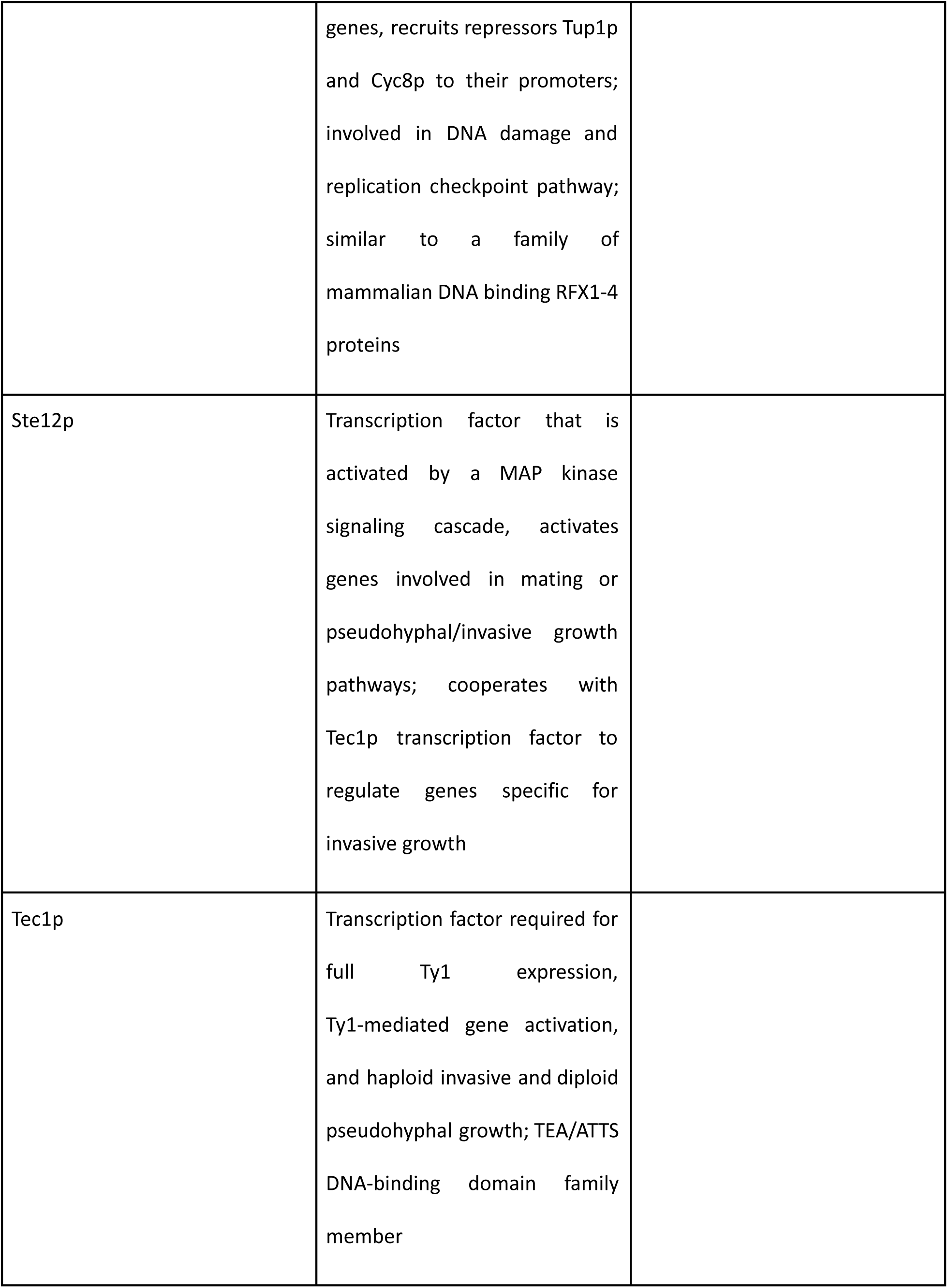
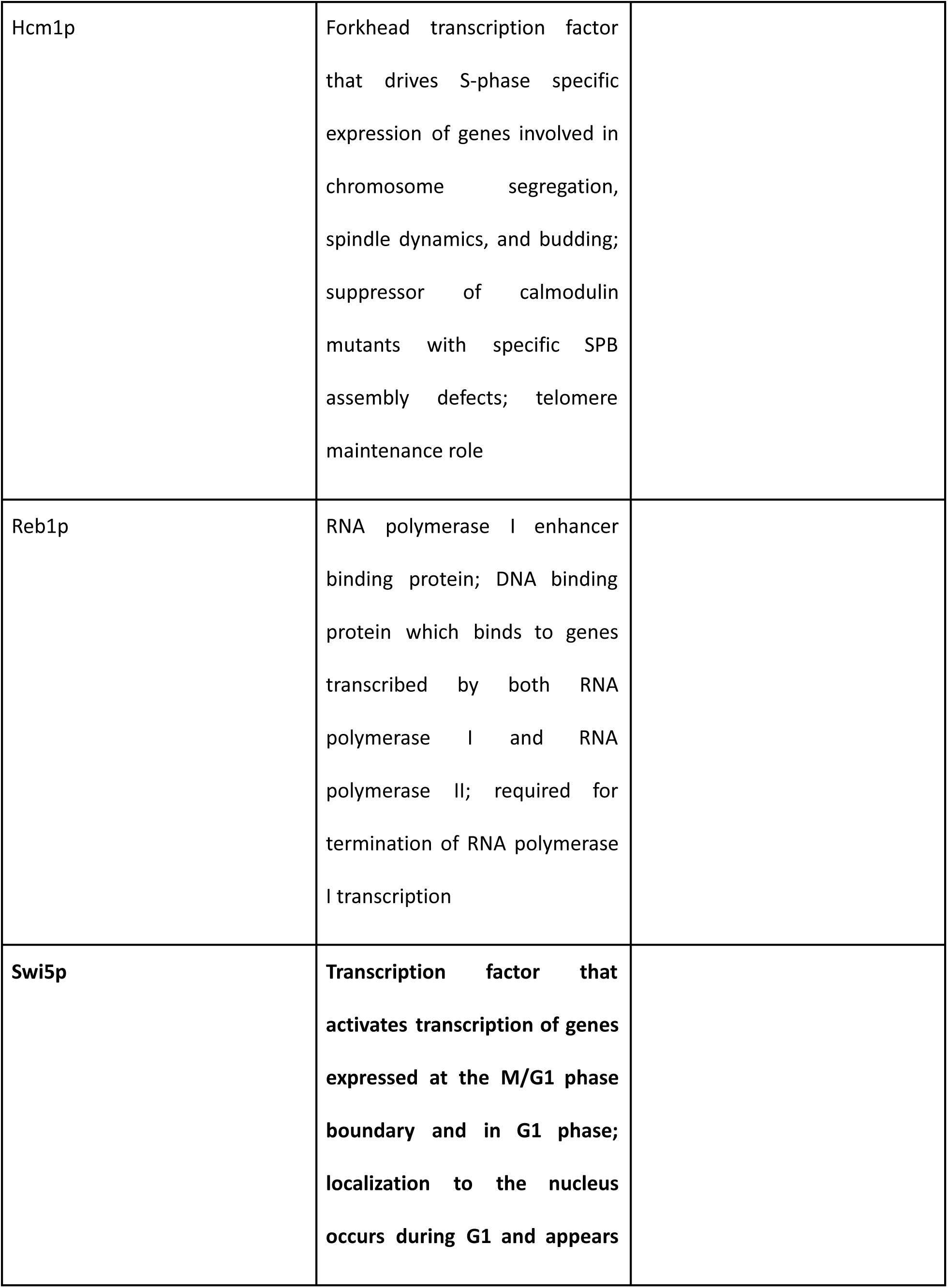
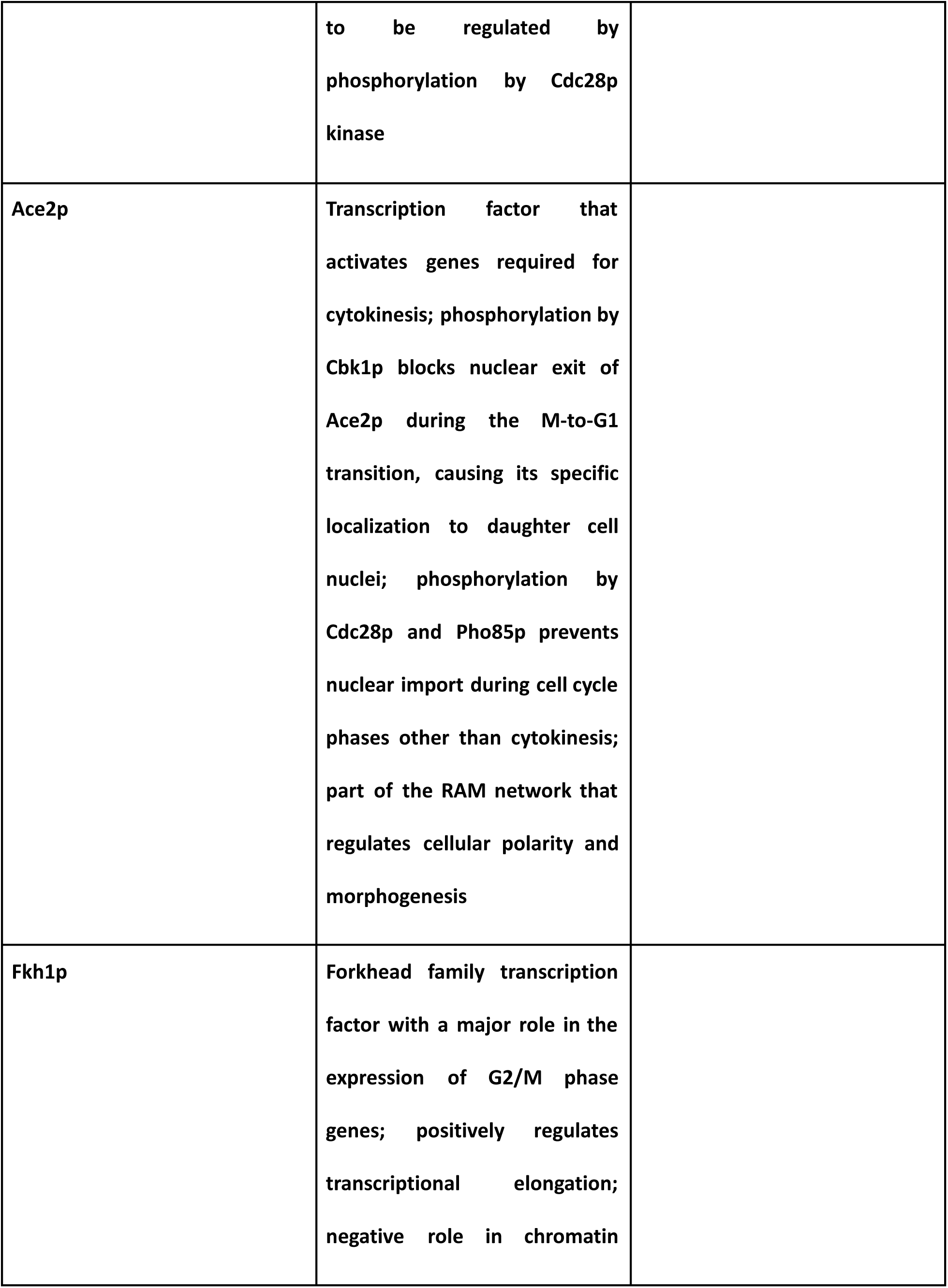
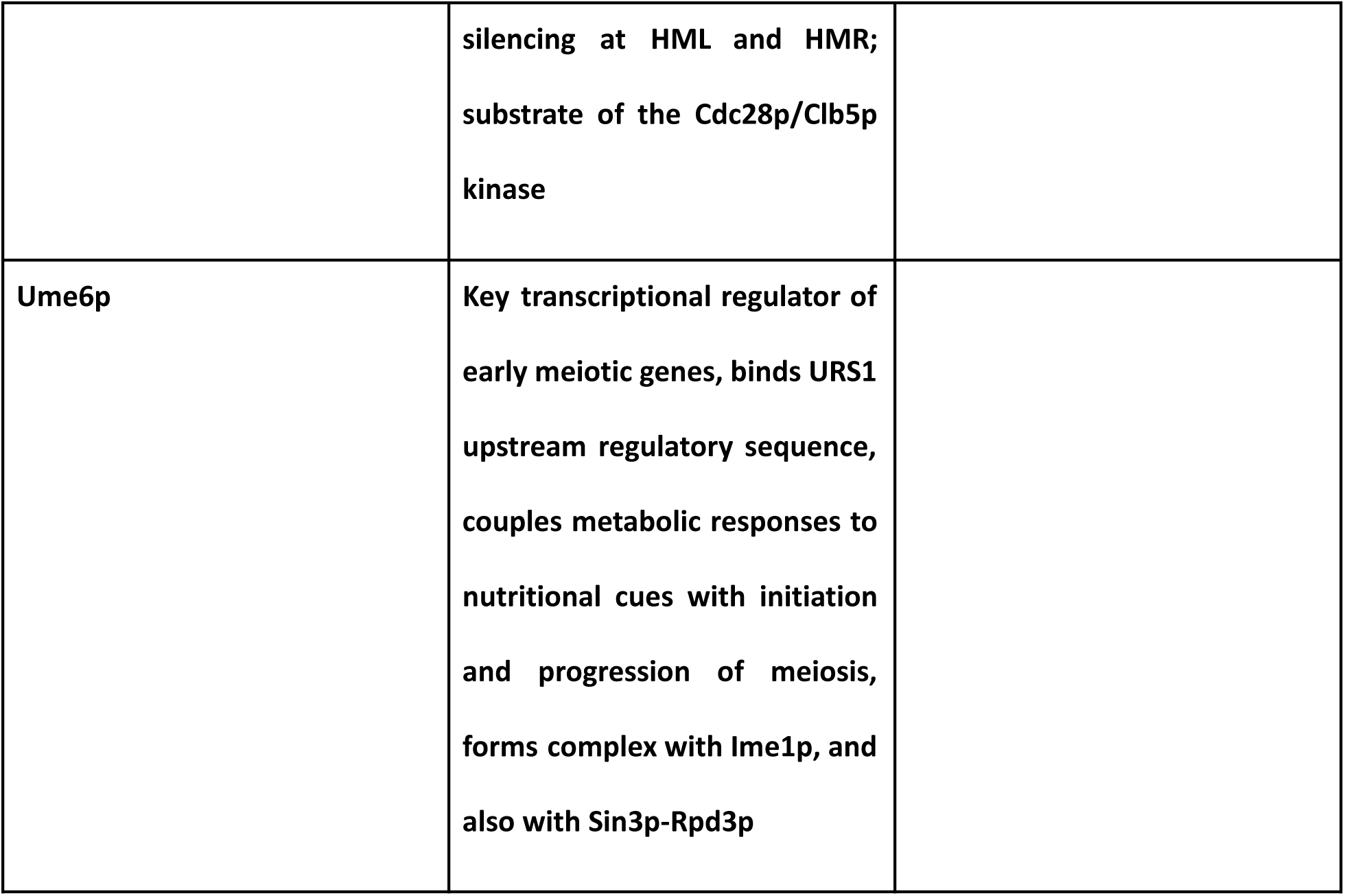

